# Microbiota-derived inosine programs protective CD8^+^ T cell responses against influenza in newborns

**DOI:** 10.1101/2024.04.09.588427

**Authors:** Joseph Stevens, Erica Culberson, Jeremy Kinder, Alicia Ramiriqui, Jerilyn Gray, Madeline Bonfield, Tzu-Yu Shao, Faris Al Gharabieh, Laura Peterson, Shelby Steinmeyer, William Zacharias, Gloria Pryhuber, Oindrila Paul, Shaon Sengupta, Theresa Alenghat, Sing Sing Way, Hitesh Deshmukh

## Abstract

The immunological defects causing susceptibility to severe viral respiratory infections due to early-life dysbiosis remain ill-defined. Here, we show that influenza virus susceptibility in dysbiotic infant mice is caused by CD8^+^ T cell hyporesponsiveness and diminished persistence as tissue-resident memory cells. We describe a previously unknown role for nuclear factor interleukin 3 (NFIL3) in repression of memory differentiation of CD8^+^ T cells in dysbiotic mice involving epigenetic regulation of T cell factor 1 (TCF 1) expression. Pulmonary CD8^+^ T cells from dysbiotic human infants share these transcriptional signatures and functional phenotypes. Mechanistically, intestinal inosine was reduced in dysbiotic human infants and newborn mice, and inosine replacement reversed epigenetic dysregulation of *Tcf7* and increased memory differentiation and responsiveness of pulmonary CD8^+^ T cells. Our data unveils new developmental layers controlling immune cell activation and identifies microbial metabolites that may be used therapeutically in the future to protect at-risk newborns.

## Main Text

Infancy is a remarkable period of vulnerability as the immune system is educated to new pathogens, particularly those infecting the lungs(*1*). Although adaptive immunity during infancy has traditionally been viewed as “immature,” recent studies suggest that infant T cells exhibit unique functional responses compared with adult T cells (*2–6*). These differences in the origin, post-thymic maturation, and homeostatic proliferation sculpt the peripheral T cell compartment, resulting in dramatic changes in the composition of the naïve T cell pool at various stages of life (*7*). A newer model for immune ontogeny, termed “layered development,” challenges the established dogma of a more linear development of the immune system from fetal life to adulthood (*8*).

Infancy also represents a critical window when diverse microbial communities are assembled in the gut(*9*). Signals from evolving microbial communities and their products can permanently program the infant’s immune system (*10, 11*), including CD8^+^ T cells (*12*). Perturbations in the composition and maturation of the intestinal microbiota in infants (*13–19*) caused by commonly used antibiotics (ABX) (*20–23*) are associated with severe viral lower respiratory tract infection (LRTI)(*24–27*) and the development of chronic inflammatory diseases during infancy (*28–30*). Whether increased short- and long-term risk to viral pathogens reflects environmental alterations to the developmental layering in the pulmonary T cell compartment remains unclear (*31*). The mechanistic basis of the developmental reprogramming of T cell responses remains unresolved.

### Decreased virus-specific CD8^+^ effector T cells and diminished persistence of tissue-resident memory cells promote influenza susceptibility in dysbiotic infant mice

To model early life ABX exposure, we treated pregnant mouse dams with ampicillin, gentamicin, and vancomycin, the most commonly used ABX in pregnant women and human newborns (*32*), beginning at embryonic day 15 (E15) and cross-fostered the pups with no-ABX treated dams at postnatal day 5 (PN5) (**Fig. 1A**). Stool of infant mice from ABX-treated dams, referred to hereafter as “dysbiotic” infant mice, demonstrated significantly less phylogenetic diversity than that of infant mice from no-ABX-treated dams, referred to hereafter as “control” (**fig. S1A-D**). Infant mice were challenged with influenza A H1N1 strain-PR8, expressing OVA_257–264_ epitope (*33*) [referred hereafter as PR8-OVA] at PN14, when newborn mouse T cells more closely approximate human infant T cells (*34*).

**Fig. 1.**
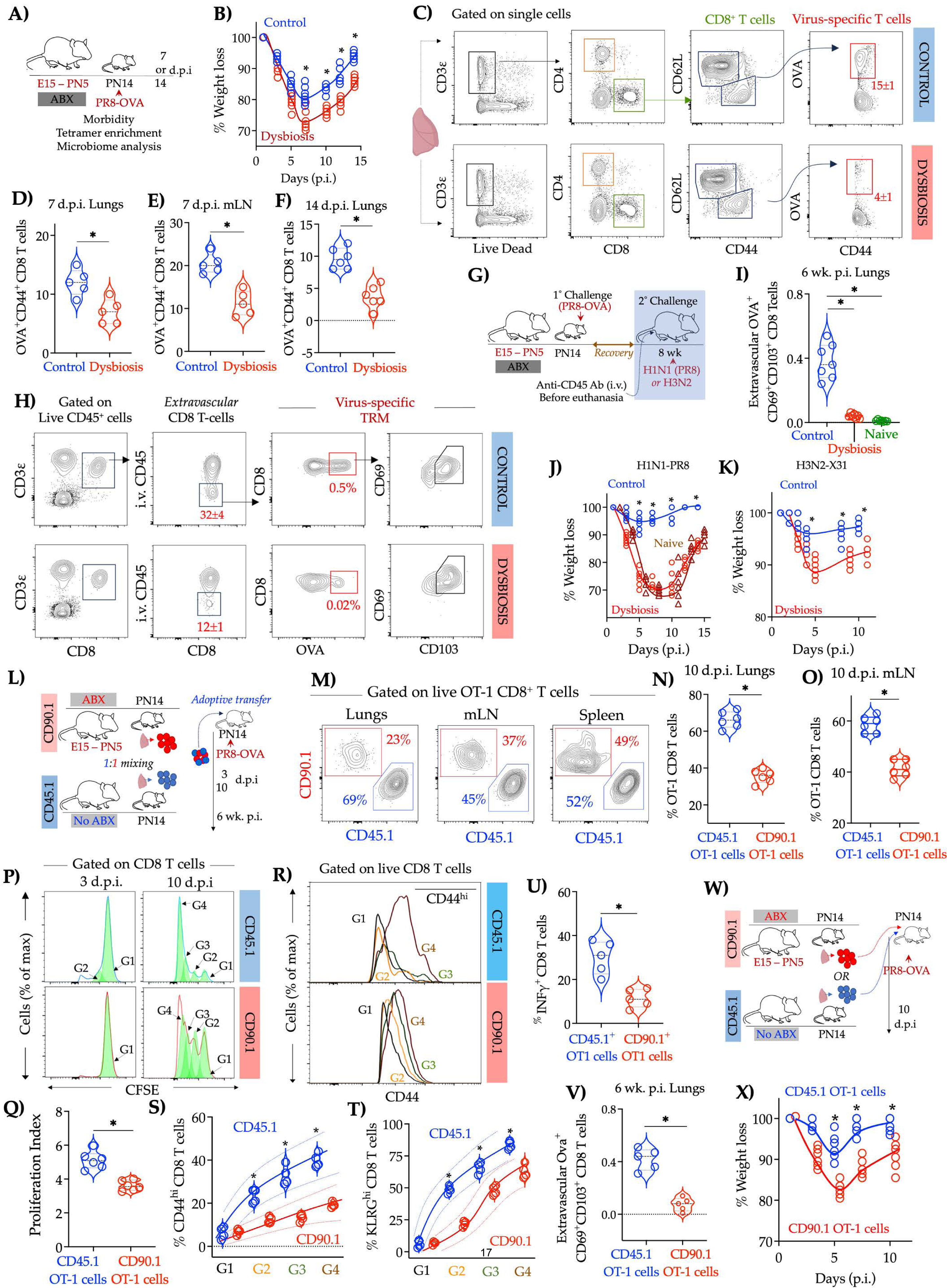
Infant mice exposed to perinatal ABX mount less robust lung localized CD8+ T cell responses to influenza A. **A)** Experimental approach. Pregnant C57/B6 dams were treated with a cocktail of antimicrobials from embryonic day (E) 15 to postnatal day (PN) 5 (dysbiosis) or with saline (control). Infant mice in each experimental group were challenged with a sublethal dose of murine-adapted influenza A H1N1 strain-PR8, expressing OVA_257–264_ epitope (PR8-OVA) [10^2^ TCID50] via the intranasal (i.n.) route on PN14. **B**) Weight change (n=6, * *p*-values < 0.05, one-way ANOVA with Tukey’s correction for multiple comparisons. Solid lines, mean; dotted lines, quartiles). **C**) Flow cytometry analysis of single-cell lung suspension from infant mice ten days post-infection (d.p.i.). Representative bi-axial plots. The proportion of OVA_257–264_ specific CD8^+^ T cells (mean ± SEM) is indicated. **D**) Proportion of OVA_257–264_ specific (tetramer^+^) CD8^+^ T cells in lungs or **E**) mediastinal lymph nodes at 7 d.p.i or **F**) lungs at 14 d.p.i. [n=5-6, * indicated *p*-values < 0.05, Student’s t-test, Solid lines, mean; dotted lines, quartiles]. **G**) Infant mice in each experimental group previously challenged with PR8-OVA were allowed to recover for six weeks and challenged with either PR8-OVA [10^3^ TCID50] or heterotypic, influenza A-strain H3N2 [10^4^ TCID50] or sham (uninfected) via i.n. route. In some experiments, age-matched mice not previously challenged with influenza during infancy (Naïve) were used. Animals were treated with fluorescent-labeled anti-CD45 antibody (Ab) via the intravenous route (i.v.) at 10 d.p.i. Thirty minutes after i.v. injection, animals were euthanized, and single-cell lung suspension was analyzed by flow cytometry **H**) Representative bi-axial plots. The proportion of extravascular or intravascular OVA_257–264_ specific CD8^+^ T cells (mean ± SEM) is indicated. **I**) Proportion of extravascular OVA_257–264_ specific CD8^+^ T cells in indicated experimental groups. [n = 7, * indicated *p*-values < 0.05, Student’s t-test, Solid lines, mean; dotted lines, quartiles]. **J**) Weight change after challenge with PR8-OVA or **K**) H3N2 (n=4-5, * *p*-values < 0.05, one-way ANOVA with Tukey’s correction for multiple comparisons. Solid lines, mean; dotted lines, quartiles). **L)** Experimental Approach. Pregnant OT-1 transgenic dams, on either CD90.1 or CD45.1 background, were treated with a cocktail of antimicrobials as described in Fig. 1A. Equal numbers of OT-1 CD8^+^ T cells from control (CD45.1) or dysbiotic (CD90.1) donors (PN14) were labeled with CFSE and adoptively co-transferred into age-matched recipient mice. Twelve hours later, the recipient mice were challenged with influenza [PR8-OVA, 10^2^ TCID50]. **M**) Flow cytometry analysis of single-cell suspension from either lung, mediastinal lymph nodes, or spleen infant mice seven days post-infection (d.p.i.). Representative bi-axial plots. The proportion of OT-1 CD8^+^ T cells from either control (CD45.1) or dysbiotic (CD90.1) host infant mice (mean) is indicated. **N**) Proportion of adoptively transferred OT-1 CD8^+^ T cells in lungs or **O**) mediastinal lymph nodes of recipient infant mice at 10 d.p.i [Solid lines, mean; dotted lines, quartiles, n=5, * indicated *p*-values < 0.05, Student’s t-test]. **P**) Representative histogram of computationally modeled CFSE peaks corresponding to cell divisions in OT-1 CD8^+^ T cells at indicated times post-infection and **Q**) proliferation index of control (CD45.1) or dysbiotic (CD90.1) OT-1 cells in recipient mouse at 10 d.p.i. (Solid lines, mean; dotted lines, quartiles, n=6, *indicated *p*-values < 0.05, one-way ANOVA with Tukey’s correction for multiple comparisons). **R**) Representative histogram of CD44 expression in OT-1 CD8^+^ T cells and **S**) Proportion of CD44^hi^ or **T**) KLRG^hi^ control (CD45.1) or dysbiotic (CD90.1) OT-1 cells in individual recipient mice at indicated cell divisions. [Solid lines, mean; dotted lines, quartiles, n=5, *indicated *p*-values < 0.05, one-way ANOVA with Tukey’s correction for multiple comparisons]. **U**) Proportion of INF γ ^+^ cells or **V)** extravascular OVA_257– 264_ specific control (CD45.1) or dysbiotic (CD90.1) TRM cells [n=6, * *p*-values < 0.05, Student’s t-test. Solid lines, mean; dotted lines, quartiles]. **W)** Infant B6 mice were adoptively transferred with OT-1 CD8^+^ T cells from age-matched control (CD45.1) or dysbiotic (CD90.1) infants on PN14. Twelve hours later, the recipient mice were challenged with influenza [PR8-OVA, 10^2^ TCID50]. **X)** Weight change was determined at indicated time points [n=6, * *p*-values < 0.05, one-way ANOVA with Tukey’s correction for multiple comparisons. Solid lines, mean; dotted lines, quartiles].

Perinatal ABX exposure did not affect the proportion of hematopoietic stem cells (HSC) (Lin^-^c-Kit^+^ cells) or common lymphoid progenitor cells (CLP) (Lin-Flt3^+^c-Kit^mid^Sca-1^mid−^CD48^+^ cells) (**fig. S1E: Gating Strategy, fig. S1 F,G**) in the bone marrow. The proportion of bulk T cells and naïve (CD44^lo^CD62L^hi^) CD8^+^ T cells was comparable in the spleen of control and dysbiotic infants (**fig. S1H: Gating Strategy, fig. S1 I-K**) before infection. After infection, the proportion of influenza-specific (CD44^+^CD62L^lo^Ova^+^) CD8^+^ T effector cells were consistently reduced in lungs and draining mLN from dysbiotic infant mice (**Fig. 1 C-F, fig. S2B: Gating Strategy)**, despite similar viral burden (**fig. S2 E**). In contrast, the proportion of bulk T cells was comparable in control and dysbiotic infant spleen and lungs after infection (**fig. S2 C,D**). Consistent with a decrease in virus-specific CD8^+^ T cells, dysbiotic mice demonstrated greater lung damage (**fig. S2 F, G**), increased mortality **(fig. S2 H)**, and weight loss (**Fig. 1B)** compared to control infant mice.

After resolution of acute infection, influenza-specific CD8^+^ T cells establish residence in the lungs, playing an essential role in protection from recurrent viral LRTI (*35, 36*). Six weeks post-infection, nearly a third of influenza-specific CD8^+^ T cells in adult mice not exposed to ABX during infancy (control) were extravascular, expressing CD69 and CD103, identifying them as CD8^+^ tissue-resident memory (TRM) (*37, 38*) (**Fig. 1 G,H**). In contrast, the proportion of influenza-specific TRM in adult mice exposed to ABX during infancy was significantly decreased (**Fig. 1 H,I**). Adult mice exposed to ABX during infancy showed more weight loss after secondary OVA-PR8 (**Fig. 1J**) or heterosubtypic H3N2 viral strain X31 (**Fig. 1K**) challenge than control mice. Weight loss and viral burden in adult mice exposed to ABX during infancy were comparable to adult mice without prior influenza A exposure (naïve adult) (**Fig. 1J**).

We treated mice with the sphingosine 1-phosphate receptor-1 agonist FTY720, which depletes circulating T cells(*39*), demonstrating that circulating T cells had a marginal role in protecting adult mice exposed to ABX during infancy after secondary OVA-PR8 challenge (**fig. S3 A,B**). CD8^+^ T cells in neonatal mice have a cell-intrinsic predisposition to exhibit a memory phenotype (Tvm) without overt immunization or infection (*40*). This fate bias has been proposed to explain the differences in proliferative capacity and diminished potential to generate long-lived memory T cells compared to adult T cells (*6, 31*). The proportion of T_VM_ remained unchanged between control and dysbiotic infants (**fig. S1K**), suggesting that diversity in the T cell pool does not entirely explain the disparities in differentiation and memory T-cell formation in dysbiotic infants.

Taken together, these data suggest that reduced proportion of influenza-specific CD8^+^ effector T cells and decreased persistence of TRM contribute to increased influenza susceptibility in dysbiotic infant mice, consistent with epidemiological data demonstrating increased viral LRTI severity and increased need for hospitalizations in infants exposed to ABX during perinatal and neonatal period (*24–27*).

### Cell-intrinsic deficits drive the divergent T cell fate post-infection in dysbiotic infant mice

Naive T cells are proposed to have identical potentials to differentiate into effector or memory T cells, their fates resolved by stochastic events following microbial infection(*41, 42*). For example, individual T cell precursors from OT-I mice, which express an identical T cell receptor (TCR), display a broad spectrum of effector phenotypes and clonal burst sizes after infection (*41*). Based on these findings, it has been speculated that the short-term and long-term fates of naive T cells could be explained by the amount and type of stimulation they experienced during infection(*43–45*). We used a co-transfer model, wherein control and dysbiotic OT-I T cells recognizing an antigenic epitope within a recombinant influenza strain (OVA)(*35, 36*) were used to compare antigen-specific responses to infection intrinsic to control or dysbiotic CD8^+^ T cells (**Fig. 1L**).

Seven d.p.i, the OT-1 cells from control infant mice outnumbered those from dysbiotic infant mice by ∼ four-fold in the lungs and, to a lesser extent, in the mLN (**Fig. 1 M-O**). CD8^+^ T cells proliferate and differentiate after antigen stimulation. Surviving antigen-specific T cells constitute the bulk of the TRM. OT-1 CD8^+^ T cells from control infant mice (CD45.1^+^ OT1) divided faster, underwent more cell divisions (**Fig. 1 P, Q and fig. S4A, B**), differentiated more rapidly, as quantified by an increased proportion of KLRG1^hi^ cells or CD44^hi^ cells per division (**Fig. 1R-T and fig. S4C**) and produced more IFNγ (**Fig. 1U** and **fig. S4D**) compared to dysbiotic infant mice (CD90.1^+^ OT1). Six weeks p.i., OT-1 CD8^+^ T from control infant mice (CD45.1^+^ OT1) generated a significantly higher number of TRM (**Fig. 1V**) compared to dysbiotic infant mice (CD90.1^+^ OT1). To assess whether cell-intrinsic deficits in CD8^+^ T-cells contributed to increased morbidity, infant mice were adoptively transferred OT-I CD8^+^ T cells from dysbiotic host, demonstrating increased weight loss and impaired recovery of body weight after infection with OVA-PR8 compared to mice adoptively transferred with OT-I CD8^+^ T cells from control infant mice (**Fig. 1W, X**). These data suggest that dysbiosis causes cell-intrinsic deficits in proliferation, differentiation of CD8^+^ effector T cells, and decreased persistence of TRM, likely contributing to increased influenza susceptibility in dysbiotic infant mice.

### Transcription factor, nuclear factor interleukin 3 regulated (NFIL3) restrains differentiation of CD8^+^ T cells in dysbiotic infant mice

We used single-cell RNA-sequencing and analysis (scRNA-Seq) to predict gene regulatory networks influenced by dysbiosis. Use of curated gene signatures enabled annotation of naive-like CD8^+^ T cells [co-expressing *Cd3d,Cd3e, Cd8a*, *Il7r* and *Sell* (encoding CD62L) and lacking activation features such as *Pdcd1* (encoding PD1) and *Tnfrsf9*] [C1]; a CD8^+^ terminally differentiated effector cluster, [high expression of *Gzmk, Gzmm, Klrd1, Ly6c* and multiple inhibitory receptors (*Pdcd1*, *Ctla4*, *Lag3*, *Tigit*, *Havcr2* (encoding TIM3) and *Tox)* [C0]; and a cell cluster with moderate to low co-expression *Tcf7 (*encoding TCF1), *Gzmk*, *Gzmm*, *Pdcd1* and *Sidt1* identified as memory CD8^+^ T cells [C3]. These cells expressed markers associated with lung residency, for example, *Cxcr6*, *Cd44, S1pr1,* and several integrins (*Itga4*, *Itgb7*, *Itgb1*). A distinct cell cluster marked by moderate expression of *Slamf6, ‘Id3*’, ‘*Tcf7*’, ‘*Pdcd1*’ and high expression of proliferation markers *Mki67* and *Top2a* approximated the transcriptomic signature of recently described proliferation-competent T cells with memory potential [C2](*46*) (**Fig. 2A-C, fig. S5A**). Monocle was used to predict relationships among T cell states, supporting our cell state assignments and revealing the temporal expression patterns of key regulators of effector differentiation (*Tcf7, Ctla4*), memory formation, and tissue residency (*Il7r* and *Sell*), (**Fig. 2D-E**). Proportion of terminally differentiated CD8^+^ effector T cells [C0] was reduced in dysbiotic infant lung (**Fig 2F**). Spectral cytometry identified analogous CD8^+^ T cell clusters (**Fig. 2G, H and fig. S5B, C**) and confirmed the remodeling of pulmonary CD8^+^ T cell pool with reduced proportion of effector and memory CD8^+^ T cells in dysbiotic infant lung (**Fig. 2I**).

**Fig. 2:**
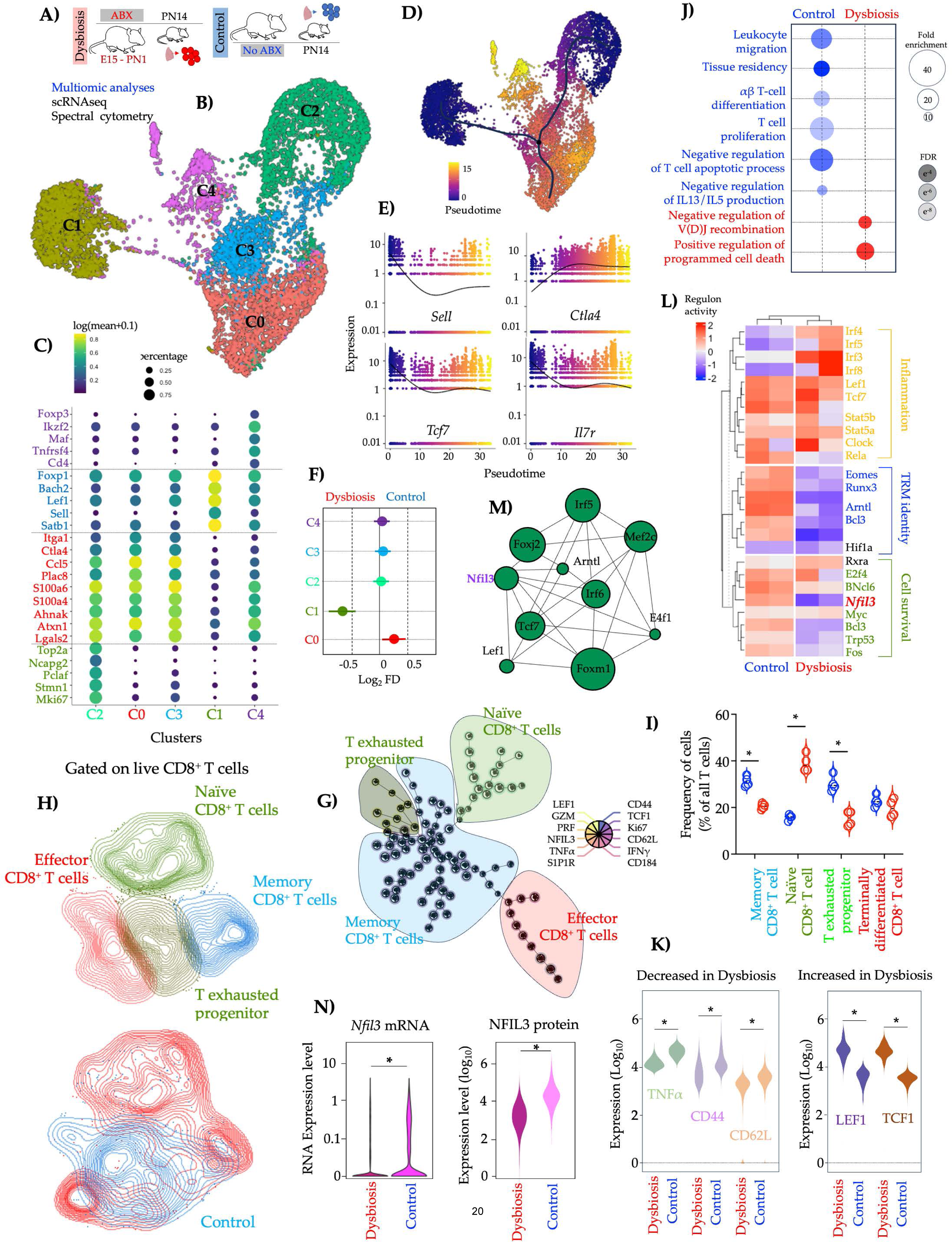
Cell-intrinsic defects in the generation of lung-localized effector T cells in dysbiotic infants. **A)** Lung from control and dysbiotic infant mice (n=4, 2 in each group) was obtained at 7 days post-infection (d.p.i). Lung samples were dissociated into cell suspensions, magnetically enriched (EPCAM^-^CD31^-^CD45^+^CD8^+^) and used for single-cell RNA sequencing (scRNAseq). **B**) Uniform manifold approximation and projection (UMAP) embedding of all samples (n∼9000 cells) colored by cell clusters was performed on scRNAseq data. C0: terminally differentiated effector CD8^+^ T cells, C1: naive-like CD8^+^ T cells C2: CD8^+^ T exhausted progenitor characterized. C3: effector-memory CD8^+^ T cells. C4: regulatory T cells. **C**) Scaled expression of genes unique to each cluster. Bubble size represents the percentage of cells expressing the gene, and the color intensity represents the log (mean expression +0.1) [Benjamini and Hochberg-adjusted *p*-values < 0.01, log_2_ fold change > 2, Wald’s test]. **D**) UMAP embedding of all samples colored by pseudo time with overlaid trajectory and **E**) scatter plots showing expression of *Sell*, *Ctla4*, *Tcf7* and *Il7r* across pseudo time. **F**) Proportion of clusters in dysbiotic and control samples showing predominance of naïve-like CD8^+^ T cells (C1) in dysbiotic mice and terminally differentiated effector CD8^+^ T cells (C0) in controls. Lines indicate 95% confidence intervals. **G)** Unsupervised analysis of live, single live CD8 T^+^ cells using a self-organizing map (SOM). CD8^+^ T cell clusters identified by SOM were mapped to **H**) uniform manifold approximation and projection (UMAP) embedding and colored by naïve, effector, memory, and exhausted progenitor clusters. **I**) The proportion of memory or naïve or exhausted progenitor or effector CD8^+^ T cells in control and dysbiotic infant lungs at ten d.p.i. [n=6, * indicated *p*-values < 0.05, NS – not significant. One-way ANOVA with Tukey’s correction for multiple comparisons. Solid lines, mean; dotted lines, quartiles). **J)** Gene Ontology (GO) terms associated with DEG in in control and dysbiotic mice. [Benjamini-Hochberg-corrected *p*-values < 0.05 (one-sided Fisher’s exact test) are shown and colored by gene ratio]. **K)** MFI of indicated markers in live CD8^+^ T cells from control or dysbiotic infant lungs [n=6 per group, * indicated *p*-values < 0.05, one-way ANOVA with Tukey’s correction for multiple comparisons. Solid lines, mean; dotted lines, quartiles]. **L**) Row-scaled regulon activity for all samples. k-means clustering was used to arrange clusters and regulons [Benjamin and Hochberg-adjusted p-values < 0.01]. Regulons related to TRM identity, cell survival, and inflammation are highlighted. **M**) Network diagram showing the core regulator relationship, with the circle size indicating # of regulated genes. **N**) Average *Nfil3* expression (left) or NFIL3 MFI (right) in CD8^+^ T cells from control or dysbiotic infant lung, at seven d.p.i. [Benjamini and Hochberg-adjusted *p*-values < 0.01].

Gene pathways associated with CD8^+^ T cell effector function and memory formation [*Cd44*, *Foxp1*, *Il4ra, Lgals1, Lgals3, Klrd1* and *Klrg1*] and cell proliferation [*Mki67* and *Top2a*] were decreased in CD8^+^ T cells from dysbiotic infant lungs (**Fig. 2J and fig. S5D**). Transcription factor (TF), *Zeb2* and its target genes *Sell* and *S1pr1* (both markers of tissue emigrating T cells), and several integrins, *Itga4* and *Itgb1* (markers of tissue residency) were decreased in CD8^+^ T cells from dysbiotic infant lungs. In orthogonal studies using spectral cytometry, we found lung CD8^+^ T cells in dysbiotic infants expressed lower levels of effector proteins, perforin 1, granzyme B, and TNFα (**Fig. 2K and fig. S5B**). The expression of tissue residency markers, CD62L and S1PR1, was lower in pulmonary CD8^+^ T cells from dysbiotic infants (**Fig. 2K and fig. S5B**). TCF1 and LEF1 are progressively downregulated during effector differentiation (*47, 48*). Low TCF1 and LEF1 protein expression levels are essential for lung TRM formation and maintenance(*49*). Consistent with these observations, we found higher TCF1 and LEF1 protein expression in lung CD8^+^ T cells from dysbiotic infants (**Fig. 2K**). Present findings suggest a remodeling of lung CD8^+^ T cell pool with a relative lack of memory-like and effector T cell signatures in CD8^+^ T cells in the lungs of dysbiotic infant mice.

SCENIC identified three distinct gene regulatory network (GRN) groups corresponding to the CD8^+^ T cell effector function, tissue residency, memory, and T cell proliferation (**Fig. 2L)**. TCF1, LEF1, and FOX01, known to regulate memory CD8^+^ T cell differentiation(*50–52*), were uniquely enriched in CD8^+^ T from control infants (**Fig. 2L**). Novel transcription factors (TF)s, including NFIL3, were inferred in GRN associated with remodeling of the CD8^+^ T pool (**Fig. 2L**). An orthogonal tool, the Inferelator 3.0, also recapitulated regulatory relationships between core TFs directing effector function, tissue residency, and cell proliferation (**fig. S5E**) and identified NFIL3 as a part of critical regulatory node directing T cell development (**Fig. 2M and fig. S5E**). NFIL3 is critical in the development and maturation of ILCs(*53, 54*) and NK cells (*55*). *Nfil3* transcripts and NFIL3 protein expression were decreased in CD8^+^ T cells in the lungs of dysbiotic infant mice (**Fig. 2N**). Therefore, we posited that NFIL3 may play a critical role in the development and maintenance of influenza-specific CD8^+^ T cells in the lungs of control infant mice.

### Decreased virus-specific CD8^+^ effector T cells and diminished persistence of TRM in mice with T cell-specific deletion of NFIL3

To generate mice that lack NFIL3 in T cells that have emigrated from the thymus after positive selection, we bred *dLck*^cre^ mice with *Nfil3*^flox^ mice(*56*) to generate *dLck*^ΔNfil3^ mice (**Fig. 3A**). Proportion of CLP and HSC were similar in the bone marrow of *dLck*^ΔNfil3^ infant mice and age-matched WT littermates (**fig. S6A, B**). Lung resident CD8^+^ T cell compartment was remodeled in *dLck*^ΔNfil3^ infant mice after challenge with PR8-OVA **(Fig. 3B**). Proportion of CD8^+^ T effector memory cells was significantly decreased at ten d.p.i. (**Fig. 3B,C**). TCF1 and LEF1 protein expression were increased in CD8^+^ T cells from *dLck*^ΔNfil3^ mice (**Fig. 3D**). Since low TCF1 expression is necessary for effector CD8^+^ T cell differentiation(*47, 48*), lung TRM formation and maintenance (*49*), we hypothesized that deficits in proliferation and effector differentiation of CD8^+^ T cells, contributed to the remodeling of CD8^+^ T cell compartment and increased influenza susceptibility in *dLck*^ΔNfil3^ infant mice. This notion was investigated by adoptively transferring equal numbers of CFSE-labeled CD8^+^ T cells obtained from PN14-day-old *dLck*^ΔNfil3^ or littermate control into age-matched hosts (**Fig. 3E**). *dLck*^ΔNfil3^ CD8^+^ T cells underwent fewer cell divisions (**Fig. 3F-H**) and differentiated more slowly, as quantified by an increase in the proportion of KLRG1^hi^ cells per division (**Fig. 3I-K)**.

**Fig. 3:**
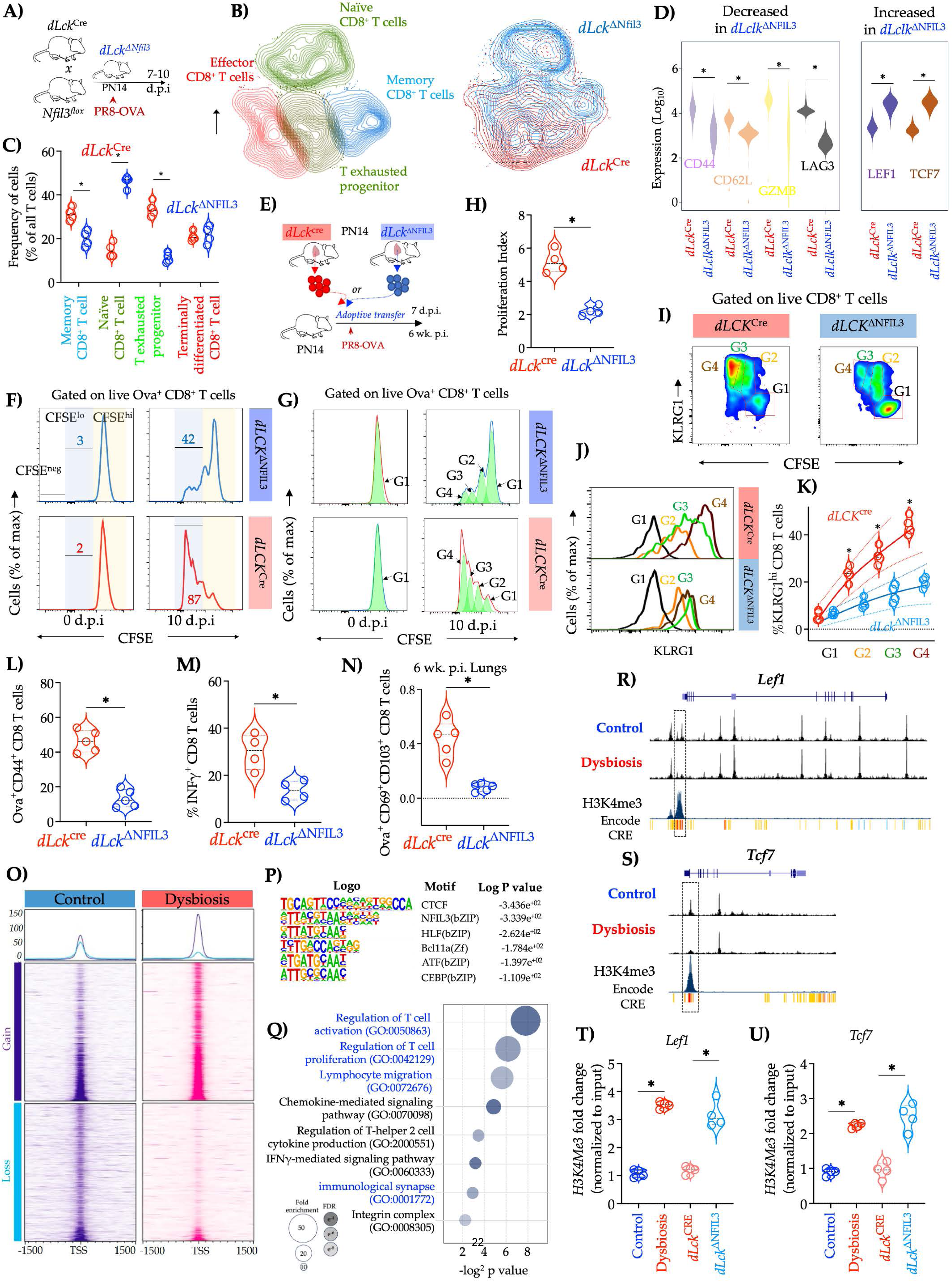
T cell-specific deletion of NFIL3 remodels the lung-localized CD8+ T cell compartment with a dominant naive T cell signature and relative lack of memory-like and effector T cells. **A)** Experimental approach. Transgenic *dLck*^cre^ and *Nfil3*^flox^ on C57Bl/6J background mice were crossbred. *dLck*^ΔNfil3^ and control littermate infant mice were challenged with a sublethal dose of murine-adapted influenza A H1N1 strain-PR8, expressing OVA_257–264_ epitope (PR8-OVA) [10^2^ TCID50] via the intranasal route on PN14. **B)** Unsupervised analysis of live, single live CD8 T^+^ cells using a self-organizing map (SOM). CD8^+^ T cell clusters identified by SOM were mapped to uniform manifold approximation and projection (UMAP) embedding and colored by naïve, effector, memory, and exhausted progenitor clusters (on left) and by genotype (right). CD8^+^ T cells from *dLck*^ΔNfil3^ infant lungs are colored blue, whereas CD8^+^ T cells from *dLck*^cre^ infant lungs are colored red. **C)** The proportion of memory or naïve or exhausted progenitor or effector CD8^+^ T cells in lungs of *dLck*^ΔNfil3^ and *dLck*^cre^ CD8^+^ T cells [n=8, * indicated *p*-values < 0.05, one-way ANOVA with Tukey’s correction for multiple comparisons. Solid lines, mean; dotted lines, quartiles]. **D)** Mean fluorescent intensity (MFI) of key phenotypic markers in live lung CD8^+^ T cells from *dLck*^ΔNfil3^ or *dLck*^cre^ infant mice [* indicated *p*-values < 0.05, one-way ANOVA with Tukey’s correction for multiple comparisons. Solid lines, mean; dotted lines, quartiles]. **E)** Experimental approach. Infant B6 mice were treated with anti-CD8α antibody (Ab) on PN12 to deplete endogenous CD8^+^ T cells and received adoptive transfer of CD8^+^ T cells from age-matched *dLck*^cre^ or *dLck*^ΔNfil3^ infants on PN14. **F**) Representative histogram of CFSE expression in CD8^+^ T cells and **G**) computationally modeled CFSE peaks corresponding to cell divisions in CD8^+^ T cells. **H**) Proliferation index of CD8^+^ T cells from lungs of *dLck*^ΔNfil3^ or *dLck*^cre^ infant mice in recipient mouse, five days after transfer [Solid lines, mean; dotted lines, quartiles, n=6, *indicated *p*-values < 0.05, Student’s t-test]. **I**) Representative biaxial plots of KLRG1 and CFSE expression in CD8^+^ T cells in recipient mice at indicated cell divisions or **J**) Representative histogram of KLRG1 expression in CD8^+^ T cells in recipient mice at indicated cell divisions. K) Proportion of KLRG1^hi^ CD8^+^ T cells from lungs of *dLck*^Δ^ ^Nfil3^ or *dLck*^cre^ infant mice recipient mouse at indicated cell divisions. [Solid lines, mean; dotted lines, quartiles, n=5, *indicated *p*-values < 0.05, one-way ANOVA with Tukey’s correction for multiple comparisons]. **L**) Proportion of OVA_257–264_ specific CD8^+^ T cells or **M)** IFNγ^+^ OVA_257–264_ specific CD8^+^ T cells or **N**) OVA_257–264_ specific TRM cells in lungs of *dLck*^ΔNfil3^ and *dLck*^cre^ infant mice [n=5, * indicated *p*-values < 0.05, Student’s t-test. Solid lines, mean; dotted lines, quartiles]. **O**) Chromatin Immuno-Precipitation with sequencing (ChiPseq) of CD8^+^ T cells isolated from lungs of control or dysbiotic infant mice at PN14 days with anti-NFIL3 antibody. Heat map of the NFIL3 ChIP-seq signals around transcription start site (TSS) compared to the random genomic control in CD8^+^ T cells from indicated infant mice. **P**) Denovo motifs identified by HOMER. **Q**) Gene Ontology (GO) terms associated with NFIL3 peaks. [Benjamini-Hochberg-corrected *p*-values < 0.05 (one-sided Fisher’s exact test) are shown and colored by gene ratio]. **R**) NFIL3 ChIP-seq or ENCODE-identified cis-regulatory elements (CRE) in *Tcf7* loci or **S**) *Lef1* loci in CD8^+^ T cells from lungs of control compared to dysbiotic mice. **T**) Fold enrichment of H3K4me3 binding at ENCODE-identified CRE in *Tcf7* or **U)** *Lef1* promoter in indicated experimental groups. [n=3, * indicated *p*-values < 0.05, one-way ANOVA with Tukey’s correction for multiple comparisons. Solid lines, mean; dotted lines, quartiles].

Influenza-specific effector CD8^+^ T cells (Ova^+^CD44^+^CD62L^lo^) (**Fig. 3L**) and IFNγ producing CD8^+^ T cells were reduced in *dLck*^ΔNfil3^ infant lungs at ten d.p.i. (**Fig. 3M**). The proportion of TRM was reduced in *dLck*^ΔNfil3^ mice six weeks post-infection (**Fig. 3N**), approximating the remodeling of lung CD8^+^ T cell pool with a decreased proportion of influenza-specific effector CD8^+^ T cells and diminished persistence of TRM observed in dysbiotic infant mice (**fig. S6C**). Taken together, our data support an essential role for NFIL3 in CD8^+^ T cell proliferation, effector function, and TRM persistence.

### NFIL3 repressed memory differentiation of CD8^+^ T cells in dysbiotic infant mice by epigenetic regulation of T cell factor 1 (TCF 1) and lymphoid enhancer-binding factor 1 (LEF1)

We cataloged the NFIL3 binding sites in the epigenome of lung-localized CD8^+^ T cells by chromatin immunoprecipitation and sequencing (Chipseq) (**Fig. 3O, P**) and identified NFIL3 binding sites in cis-regulatory regions (CRE)s of critical genes involved in T-cell memory and tissue residency, T cell maturation, and bioenergetics (**Fig. 3Q**). Chromatin binding of NFIL3 at CRE in *Tcf7* and *Lef1* locus was decreased in CD8^+^ T cells from dysbiotic infants (**Fig. 3R, S)**. Reciprocally, TCF1 and LEF1 protein expression was increased in lung CD8^+^ T cells from dysbiotic infants (**Fig. 3D**), raising the possibility that NFIL3 may repress *Tcf7* and *Lef1* transcription, to inhibit proliferation and effector differentiation of CD8^+^ T cells in dysbiotic infant lung.

To determine if NFIL3 bound to CRE and modified chromatin at *Tcf7* or *Lef1* locus to repress *Tcf7* and *Lef1* gene transcription, we examined tri-methylation of lysine 4 on histone H3 (H3K4me3), a histone mark associated with active gene transcription. Consistent with the hypothesis that NFIL3 represses *Tcf7* and *Lef1* transcription, CD8^+^ T cells from dysbiotic infants demonstrated decreased NFIL3 chromatin binding (**Fig. 3 R,S**) and increased H3K4me3 enrichment in the *Tcf7* or *Lef1* locus (**Fig. 3T, U**). To test whether NFIL3 is required for this repressive chromatin modification in the *Tcf7* locus, we compared H3K4me3 enrichment in lung *dLck*^ΔNfil3^ CD8^+^ T cells. The absence of NFIL3 resulted in increased H3K4me3 enrichment in the proximal promoter region of the *Tcf7* and *Lef1* and increased *Tcf7* and *Lef1* mRNA (**Fig. 3T, U, fig. S6F, G**). TCF7 and LEF1 work collaboratively to organize the genome of CD8^+^ T cells in a fashion necessary for CD8^+^ T cell function and identity (*57, 58*). Our data suggest that NFIL3 acts upstream of these two important regulators of T cell identity, repressing them to diminish proliferation and effector CD8^+^ T cell differentiation.

### Deficits in T cell proliferation, effector differentiation, and TRM abundance in the lungs of dysbiotic human infants

A major limitation in studying human immune responses is that sampling is confined mainly to peripheral blood, whereas adaptive immune responses are generated in tissues(*59*). Access to post-mortem lung tissues from infants (**Fig. 4A, table S1**). allowed us to overcome inherent challenges in studying lung-resident immune cells, such as limited access to tissues and difficulty manipulating these cells. We identified T cell types, including naïve CD4^+^ T cells, CD4^+^ T effector memory cells (CD4 TEM), CD8^+^ T effector memory cells (CD8 TEM), CD8 TRM cells using curated tissue-specific T-cell state signatures(*60, 61*) (**Fig. 4B,C**). Transcriptional changes in T cell clusters in the dysbiotic human infants were similar to those in dysbiotic infant mice (**Fig. 4D-F and fig. S7A**), with decreased expression of *IKZ2F*, *SOX4,* and *ID3* (TF implicated in T cell development), *TOP2A* and *KI67* (proliferation markers) and *SELL, CXCR4*, *ITGA1* and *ITGB1* (markers of tissue residency) in T cells from dysbiotic human infant lungs (**fig. S7 A**). Gene regulatory network analysis predicted that NFIL3 is a key regulator underlying the transcriptional remodeling of lung CD8^+^ T cells (**Fig. 4G**). These findings highlight the similarities in gene networks regulating T cell proliferation, effector differentiation, and TRM abundance in human infants and mice treated with antibiotics, underscoring the translational relevance of our work.

**Fig. 4:**
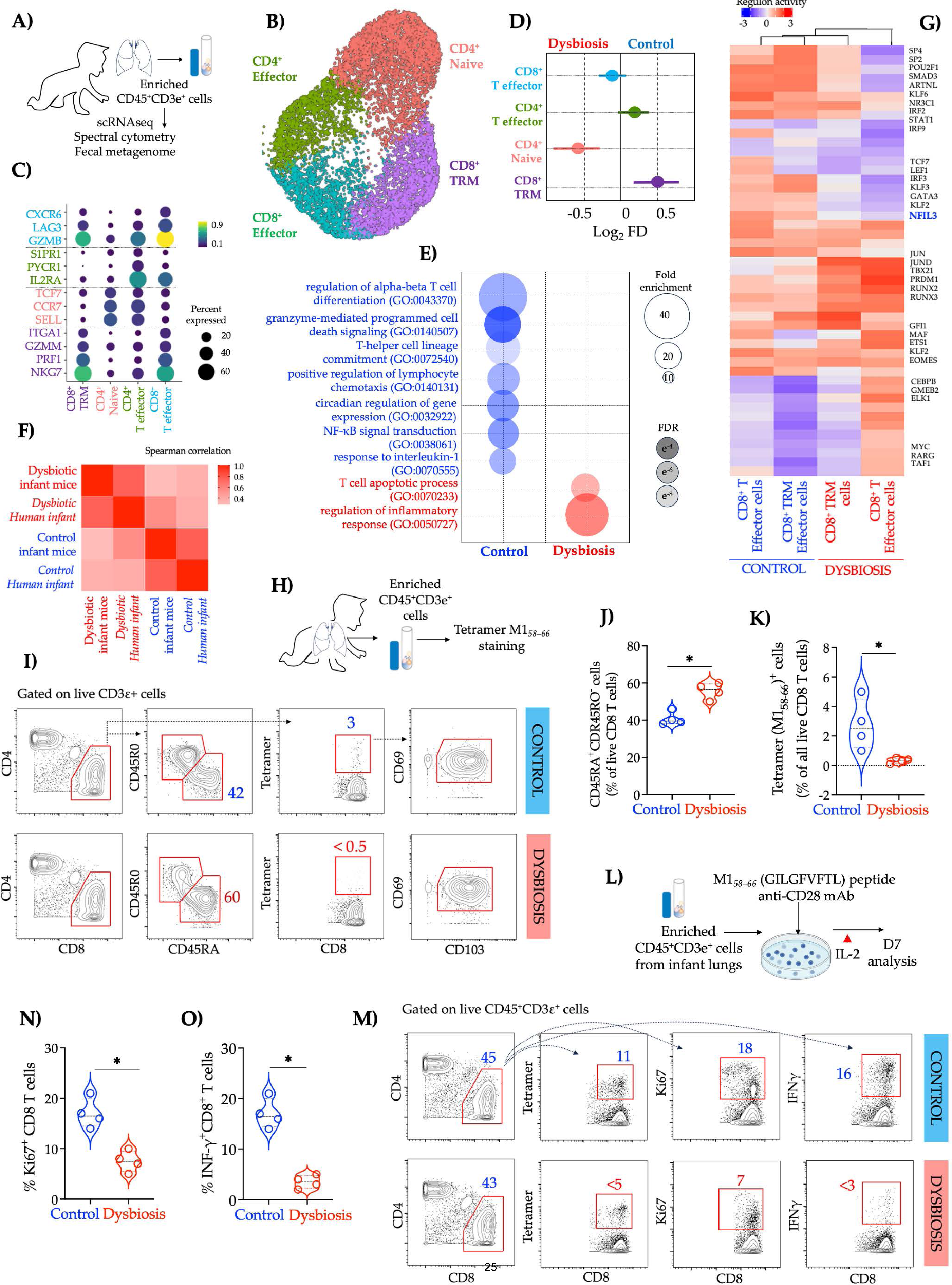
Cell-intrinsic defects in the generation of lung-localized effector T cells in dysbiotic human infants. A) Enriched (CD45^+^CD3ε ^+^) single-cell suspension from control and dysbiotic human infants (n=4, 2 in each group) were used for single-cell RNA sequencing (scRNAseq). **B**) Uniform manifold approximation and projection (UMAP) embedding of all samples (n∼6000 cells) colored by cell clusters. **C**) Scaled expression of genes unique to each cluster. Bubble size represents the percentage of cells expressing the gene, and the color intensity represents the log (mean expression +0.1) [Benjamini and Hochberg-adjusted *p*-values < 0.01, log_2_ fold change > 2, Wald’s test]. **D**) Proportion of clusters in dysbiotic and control samples showing predominance of CD8 TRM in controls. Solid lines represent 95% confidence interval. **E)** Gene Ontology (GO) terms associated with DEG in control or dysbiotic infants. [Benjamini-Hochberg-corrected *p*-values < 0.05 (one-sided Fisher’s exact test) are shown and colored by gene ratio]. **F)** Heatmap plot showing Spearman correlation of paired transcriptional profiles of control and dysbiotic human infants and control and dysbiotic infant mice. **H)** Enriched (CD45^+^CD3ε ^+^) single-cell suspension from control and dysbiotic human infants were analyzed by flow cytometry. **I)** Representative biaxial plots. Proportion of CD45RA^+^CD45RO^-^ CD8^+^ T cells or tetramer^+^ CD8^+^ T cells, co-expressing residency markers, CD103 and CD69 in indicated. **J**) Proportion of CD45RA^+^CD45RO^-^ CD8^+^ T cells or **K)** Tetramer^+^ CD69^+^CD103^+^ CD8^+^ T cells (TRM) in dysbiotic and control human infants [Solid lines, mean; dotted lines, quartiles, n=6, *indicated *p*-values < 0.05, student’s t test]. **L**) Enriched (CD45^+^CD3ε ^+^) single-cell suspension from control and dysbiotic human infants were co-incubated with >M158>–66 (GILGFVFTL) peptide and anti-CD28 mAb, followed by stimulation with IL-2 and analyzed by flow cytometry after seven days. **M)** Representative biaxial plots. Proportion of tetramer^+^ CD8^+^ T cells or CD8^+^ T cells co-expressing Ki67 or IFNγ is indicated. **N**) Proportion of KI67^+^ CD8^+^ T cells or **O**) INFγ^+^ CD8^+^ T cells in dysbiotic and control human infants [Solid lines, mean; dotted lines, quartiles, n=5, *indicated *p*-values < 0.05, Student’s t test].

We compared pre-existing influenza-specific CD8^+^ T cells using an MHCI M158– 66 (GILGFVFTL) tetramer, a highly conserved IAV epitope (**Fig. 4H**)(*62*). CD8^+^ T cells from the dysbiotic human infants exhibited predominantly naïve (CD45RA^hi^CD45R0^lo^) phenotype (**Fig. 4I, J, fig. S7B: Gating Strategy**). The proportion of influenza-specific TRM (M1*_58-66_*^+^CD69^+^CD103^+^CD8^+^ T cells) was decreased in CD8^+^ T cells from dysbiotic human infants (**Fig. 4K**). Next, we compared the functional capacity of lung CD8^+^ T cells from dysbiotic human infants to proliferate and differentiate after stimulation with cognate antigen, M1*_58–66_* peptide (**Fig. 4L**). Deficits in expansion and effector function after antigen stimulation were quantified, demonstrating a decreased proportion of proliferating CD8^+^ T cells (Ki67^+^ CD8^+^ cells) (**Fig. 4M, N**) and IFNγ producing CD8^+^ T cells (IFNγ^+^CD8^+^ cells) (**Fig. 4M, O**) in lungs from dysbiotic infants.

Our data suggest that size and quality of the CD8^+^ T-cell response to the influenza virus was inferior in dysbiotic infants. Similar attrition of lung TRM is seen in the elderly (*63*) who exhibit a diminished ability to respond to and clear respiratory infections. Based on our murine data, we speculate that depleted virus-specific TRM in dysbiotic human infants may be driven by the small size of the human lung TRM pool during infancy, compounded by an impaired ability to replenish this memory T-cell subset in dysbiosis. These data highlight how early-life microbiota exposure represents an evolutionarily conserved layer of regulation in T cell ontogeny across species.

### Microbiome-derived inosine mediates CD8^+^ T cell proliferation, effector differentiation, and increased TRM persistence

While fecal transplantation has improved clinical outcomes and CD8^+^ T cell responses in immunotherapy trials (*64, 65*) and immune imprinting in newborn mice (*66, 67*), concerns related to safety and efficacy of fecal transplantation have restrained their clinical use in vulnerable infants (*68*). The safest and most cost-effective way to promote lung immunity in infants may be by administering microbial factors or metabolites (*69*). Metagenomic sequencing of intestinal contents from control and dysbiotic infant mice and fecal specimens from control or dysbiotic human infants revealed converging trends (**Fig. 5A-C and fig. S8A, B**). *Bifidobacterium* was identified as abundant in the stools of human and murine infants (**Fig. 5B, C**). Inosine and degradation products of inosine, such as hypoxanthine, were enriched in the fecal microbiota of control human infants and newborn mice (**fig. S8C, D**). *Bifidobacterium pseudolongum* (**Fig. 5B**), abundant in the fecal microbiota of control human infants, produced inosine (*70*). Inosine and hypoxanthine were increased in the sera of *Bifidobacterium pseudolongum*-colonized mice (*70*). Consistent with the absence of *Bifidobacterium,* levels of inosine and hypoxanthine were significantly decreased in the plasma of dysbiotic mice (**fig. 5D**).

**Fig. 5:**
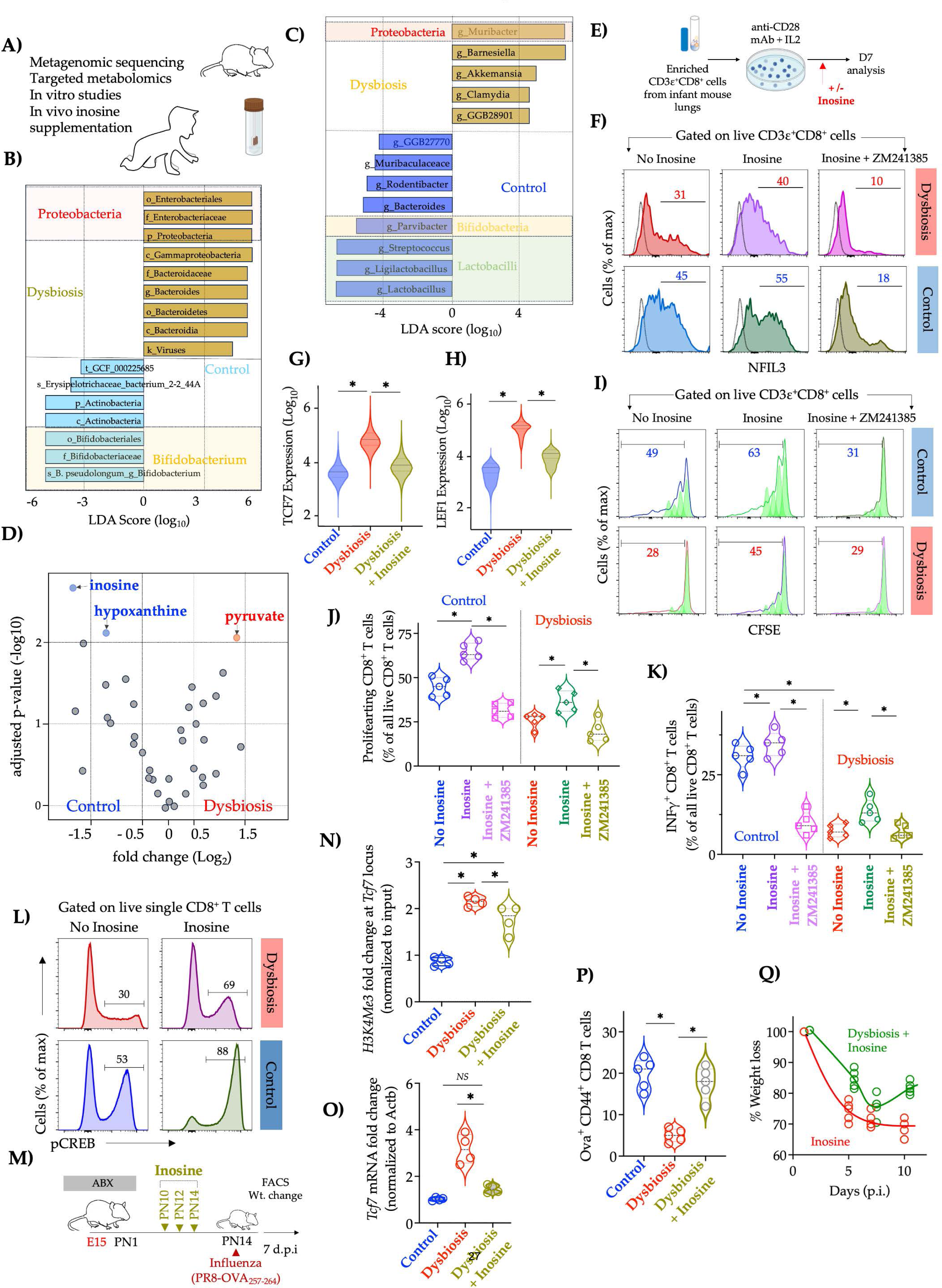
Microbiome-derived inosine is essential for robust lung-localized anti-viral response and NFIL3-directed remodeling of the lung-T cell compartment. **A)** Experimental approach. Fecal samples from control and dysbiotic humans and intestinal contents from control or dysbiotic infant mice samples were subjected to shotgun metagenomics and targeted metabolomics to quantify differences in microbiota taxonomy and function. **B)** Linear discriminant analysis effect size showing the differentially abundant clades between dysbiotic and control infant mice with an abundance of Bifidobacteria and Lactobacilli genera in controls and Proteobacteria in dysbiotic infant mice. **C)** Linear discriminant analysis effect size showing the differentially abundant clades between dysbiotic and control infant humans with an abundance of Bifidobacterium in controls and Proteobacteria in dysbiotic infant humans. **D)** Volcano plot depicting the abundance of various metabolites in plasma of dysbiotic and control infant mice. **E)** Enriched (CD3ε^+^ CD8^+^) single-cell suspension from control and dysbiotic infant mice were co-incubated with inosine with or without an A2AR antagonist (ZM241385). **F)** Representative histogram of NFIL3 expression on CD8^+^ T cells in indicated experimental groups. Proportion of NFIL3^+^ CD8^+^ T cells is indicated. **G**) Mean fluorescent intensity (MFI) of TCF1 or **G)** LEF1 in CD8^+^ T cells in indicated experimental groups. [* indicated *p*-values < 0.05, n=5, Student’s t-test. Solid lines, mean; dotted lines, quartiles]. **I**) Representative histogram of CFSE expression and **J**) proportion of proliferating CD8^+^ T cells or **K**) IFNγ^+^ CD8^+^ T cells in indicated experimental groups [n=5, * indicated *p*-values < 0.05, one-way ANOVA with Tukey’s correction for multiple comparisons. Solid lines, mean; dotted lines, quartiles]. **L**) Representative histogram of phosphorylated CREB (pCREB) expression. Proportion of pCREB^+^ CD8^+^ T cells is indicated. **L)** Experimental approach. Pregnant C57/B6 dams were treated with a cocktail of antimicrobials from embryonic day (E) 15 to postnatal day (PN) 5 (dysbiosis) or with saline (control). Infant mice in each experimental group were treated with inosine (300 μg g^-1^) on PN10,12 and 14 via intraperitoneal route and subsequently challenged with a sublethal dose of murine-adapted influenza A H1N1 strain-PR8, expressing OVA_257–264_ epitope (PR8-OVA) [10^2^ TCID50] via the intranasal (i.n.) route on PN14. **N**) Fold enrichment of H3K4me3 binding at ENCODE-identified CRE in *Tcf7* promoter in lung CD8^+^ T cells in indicated experimental groups. [n=3, * indicated *p*-values < 0.05, one-way ANOVA with Tukey’s correction for multiple comparisons. Solid lines, mean; dotted lines, quartiles]. **O)** Fold change in *Tcf7* transcripts in lung CD8^+^ T cells in indicated experimental groups. [n=3, * indicated *p*-values < 0.05, one-way ANOVA with Tukey’s correction for multiple comparisons. Solid lines, mean; dotted lines, quartiles]. **P**) Proportion of OVA_257– 264_ specific CD8^+^ T cells in lungs [n=5, * indicated *p*-values < 0.05, one-way ANOVA with Tukey’s correction for multiple comparisons. Solid lines, mean; dotted lines, quartiles] or **Q**) weight change in indicated experimental groups (n=6, * *p*-values < 0.05, one-way ANOVA with Tukey’s correction for multiple comparisons. Solid lines, mean; dotted lines, quartiles).

To test whether inosine could influence proliferation and effector maturation, CD8^+^ T cells from dysbiotic infant mice were co-incubated with or without inosine (**Fig. 5E**). NFIL3 protein expression in dysbiotic lung CD8^+^ T-cells increased after co-incubation with inosine (**Fig. 5F, fig. S9A).** Inosine suppressed TCF7 and LEF1 protein expression in dysbiotic lung CD8^+^ T-cells (**Fig. 5G, H**) and increased the numbers of both proliferating CD8^+^ T-cells (**Fig. 5I, J**) and IFNγ-producing CD8^+^ T-cells (**Fig. 5K and fig. S9B**). Similarly, co-incubation with inosine increased the proliferation (**fig. S10B, C**) and activation (**fig. S10D,E**) of CD8^+^ T cells from dysbiotic human infants. Since inosine activates adenosine 2A receptors (A2AR), to enhance cAMP-PKA signaling and phosphorylation of CREB (pCREB), a known transcriptional activator of *Nfil3* (*71*), we tested whether inosine influenced proliferation and effector maturation of CD8^+^ T cells via A2AR-pCREB pathway. In vitro, inosine increased pCREB expression in CD8^+^ T cells from dysbiotic infant mice (**Fig. 5L**). Pharmacological inhibition of A_2A_R signaling by co-incubation with the high-affinity antagonist ligand ZM241385 decreased NFIL3 protein expression (**Fig. 5F and fig. S9A**) and attenuated the increase in the proportion of proliferating CD8^+^ T-cells (**Fig. 5I, J**) and IFNγ^+^ CD8^+^ T cells from dysbiotic infant mice (**Fig. 5K and fig. S9B**).

To test whether inosine influences the proliferation and maturation of CD8^+^ T cells *in vivo*, dysbiotic infant mice were treated with inosine before influenza infection (**Fig. 5M**). H3K4me3 enrichment in the proximal promoter region of the *Tcf7* locus (**Fig. 5N**) decreased in the lung CD8^+^ T cells from dysbiotic infant mice treated with inosine, similar to H3K4me3 enrichment observed in control infant mice. Reciprocally, *Tcf7* gene transcripts were decreased in the lung CD8^+^ T cells from dysbiotic infant mice treated with inosine (**Fig. 5G, O**). Influenza-specific CD8^+^ T effector cells were increased in dysbiotic infant mice treated with inosine (**Fig. 5P**), consistent with recent observations that inosine can sustain T cell differentiation(*70, 72*). Remarkably, inosine reversed the increased morbidity of dysbiotic infant mice to influenza (**Fig. 5Q)**. Metabolic activity of the recruited cells, especially neutrophils, can lead to a local decrease in glucose levels(*73*), impacting T-cell function at sites of inflammation. We speculate that inosine is an alternative fuel for T cells in influenza-infected lungs sustaining metabolically demanding effector functions when glucose is scarce (*74*).

Together, these results highlight mechanisms behind early life CD8^+^ T cell susceptibility following dysbiosis across species and unveil new developmental layers controlling immune cell activation and identify microbial metabolites that may be used therapeutically in the future to protect infants against severe viral pneumonia (**fig. 11**). Our results challenge current views of infancy as a window of susceptibility and reimagine it as a window of opportunity to improve pulmonary health and disease burden throughout childhood.

## Acknowledgements

We thank the Cincinnati Children’s Hospital Research Foundation’s Single Cell Genomics Core and Flow Cytometry Laboratories (supported by AR47363, DK78392, and DK90971 from the National Institutes of Health [NIH]). We thank Drs. Masato Kubo and Lora Hooper for the *Nfil3*^loxp^ mice. We thank J. Whitsett, J. Chen, and C. Pasare for their helpful comments.

## Author contributions

H.D., S.S.W., J.S., and J.K. designed the experiments.

S.S.W, J.K., G.P., S.S., W.Z., and T.A. provided tools.

G.P. and L.P. provided the patient-related metadata.

H.D., J.S., E.C., W.Z., M.B., F.A., and S.S. analyzed the single-cell, bulk-transcriptomic, and metagenomic data.

J.S., J.G., and H.D. analyzed the cytometry data.

J.S., T.A., and A.R. performed experiments in mouse models.

J.S. and H.D. wrote the manuscript with inputs from S.S.W, M.B., S.S., W.Z., T.A. and S.S.

## Data

All data associated with this study are present in the paper or supplementary materials. Data discussed in this publication have been deposited in NCBI’s Sequence Read Archive (BioProject accession number PRJNA1095971). Processed scRNAseq data can be further explored at https://shelby-steinmeyer.shinyapps.io/Mouse_and_Human/

## Code availability

Scripts used for data analysis are available from GitHub at: https://github.com/Deshmukh-Lab/2024_Stevens_Influenza/.

## Resource and Reagent availability

Further information and requests for resources and reagents should be directed to and fulfilled by the Lead Contact, Hitesh Deshmukh. The *Nfil3* ^loxp^ mice used in this study were obtained under an MTA agreement from RIKEN that does not allow for the distribution of lines generated. We will provide the *Nfil3*^loxp^ mice to requesting labs that have received approval for using these mice via an MTA with RIKEN.

## Supplemental information

Figures S1–S11 and Table S1-S4

**Fig S1:**
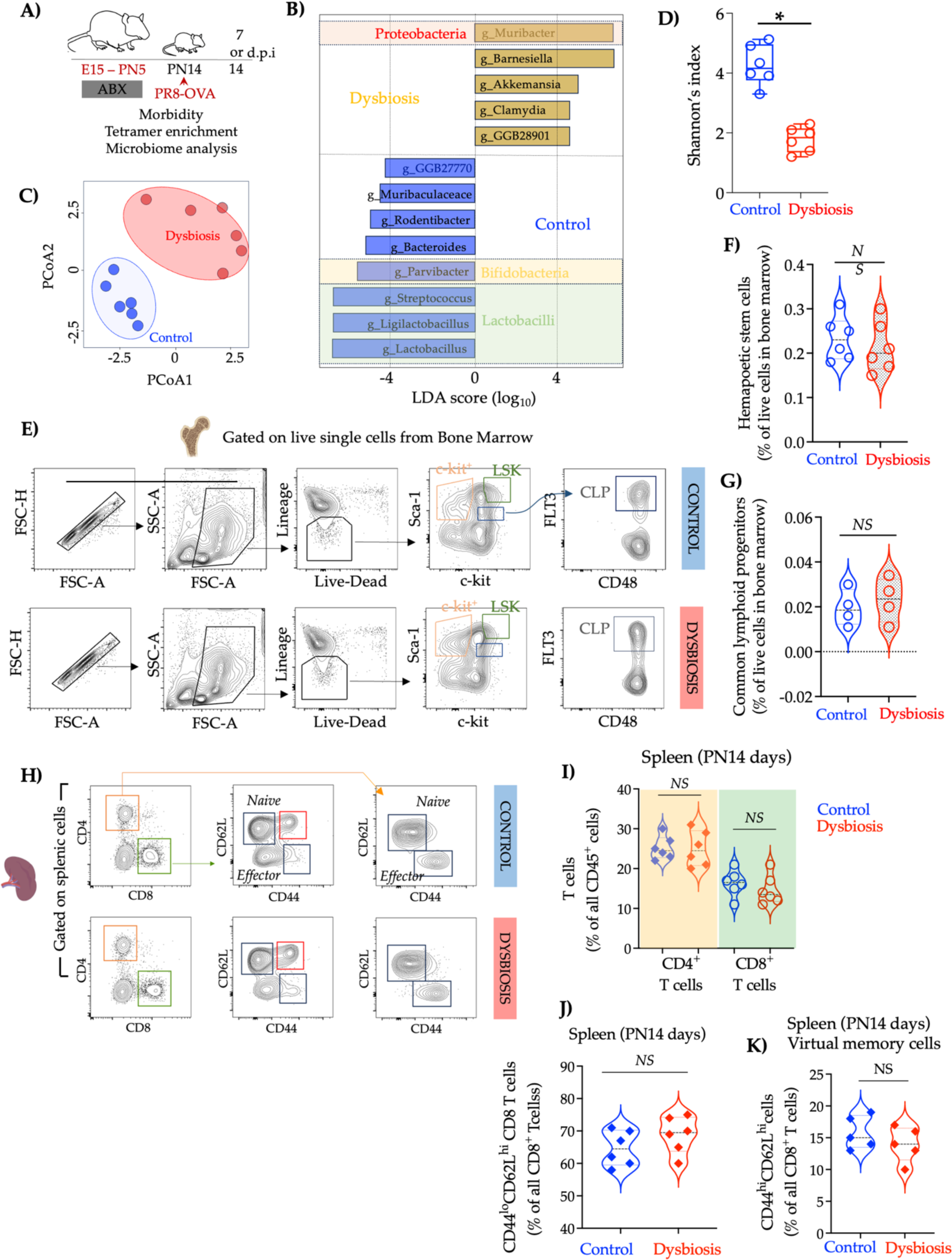
Altered gut microbiota in infant mice exposed to perinatal ABX (related to Fig. 1). **A)** Experimental approach. Pregnant C57/B6 dams were treated with a cocktail of antimicrobials from embryonic day (E) 15 to postnatal day (PN) 1 (dysbiosis) or with saline (control). Intestinal contents and bone marrow were harvested on PN14. **B)** Linear discriminant analysis effect size showing the differentially abundant clades between gut microbiota of dysbiotic and control infant mice. **C)** Principal Component (PCA) of fecal microbial communities of dysbiotic and control infant mice. **D**) β-diversity (unweighted UniFrac) of fecal bacterial communities of dysbiotic and control infant mice [n=8, * *p*-values < 0.05, Tukey bars, lines at mean and quartiles]. **E)** Flow cytometry analysis of single cell-bone marrow suspension from dysbiotic and control infant mice (PN14). Representative biaxial plots. Hematopoietic stem cells (HSC) were identified as Lin^-^c-Kit^+^ cells and common lymphoid progenitor cells (CLP) were identified as Lin-Flt3^+^c-Kit^mid^Sca-1^mid−^CD48^+^ cells. **F**) Proportion of HSC and **G**) CLP in the bone marrow of control and dysbiotic infants [n =6, Student’s t test, NS – not significant, violin plots, Solid lines, mean; dotted lines, quartiles]. Gating strategy to identify OVA257–264 specific CD8^+^ T cells in the lung. Representative flow cytometry plots. **H)** Flow cytometry analysis of single-cell suspension from spleen at PN14 prior to infection. Representative bi-axial plots. **I)** Proportion of bulk CD4^+^ or CD8^+^ T cells (expressed as a percentage of all CD45^+^ cells) or **J)** Proportion of CD44^lo^CD62L^hi^ CD8^+^ T cells or **K**) CD44^hi^CD122^hi^ CD8^+^ T cells (*virtual memory T cells*) (expressed as a percentage of all CD8^+^ T cells in the spleen. [n=5-6, NS – not significant, Student’s t-test, violin plots, Solid lines, mean; dotted lines, quartiles].

**Fig. S2:**
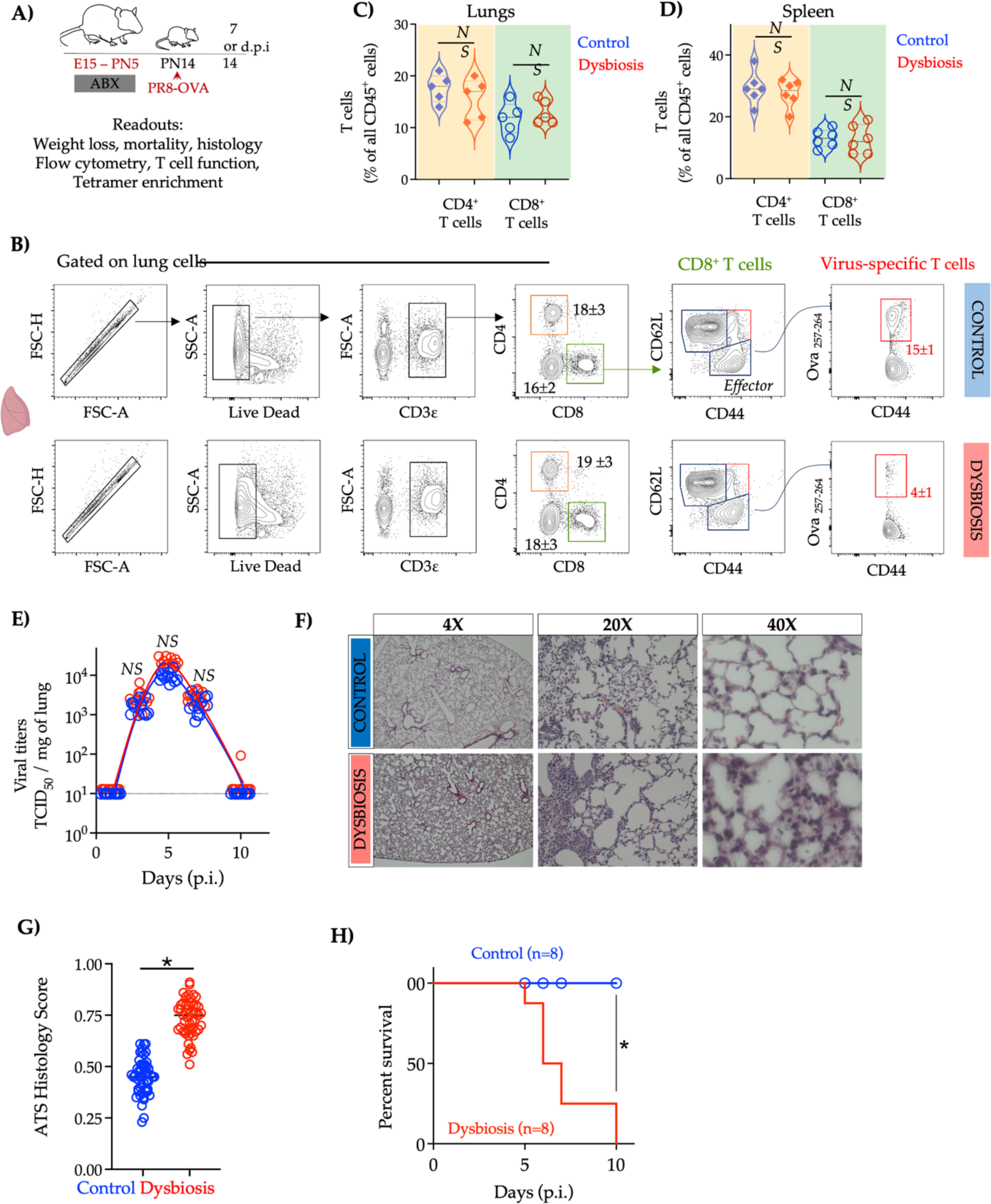
Dysbiotic infant mice exhibit greater lung damage after infectious challenge with influenza A (Related to Fig. 1). **A)** Experimental approach. Pregnant C57/B6 dams were treated with a cocktail of antimicrobials from embryonic day (E) 15 to postnatal day (PN) (dysbiosis) or with saline (control). Infant mice in each experimental group were challenged with a sublethal dose of murine-adapted influenza A H1N1 strain-PR8, expressing OVA_257–264_ epitope (PR8-OVA) [10^2^ TCID50] via the intranasal (i.n.) route on PN14. **B)** Gating strategy to identify OVA257–264 specific CD8^+^ T cells in the lung. Representative flow cytometry plots. Numbers (mean ± SEM) represent the proportion of indicated populations. **C**) Proportion of bulk CD4^+^ or CD8^+^ T cells (expressed as a percentage of all CD45^+^ cells) in the lungs or **D**) spleen 10 days post-infection (d.p.i.). [n=5-6, NS – not significant, Student’s t-test, violin plots, Solid lines, mean; dotted lines, quartiles]. **E)** The viral titers of influenza A H1N1 strain-PR8 at indicated times in the lungs of control or dysbiotic mice. The individual data points for dysbiotic mice are offset for clarity. [Solid lines, mean; dotted lines, quartiles, NS – not significant, one-way ANOVA with Tukey’s correction for multiple comparisons. Solid lines, mean; dotted lines, quartiles]. **F**) Photomicrographs of hematoxylin and eosin (H&E) stained sections of lungs from either control or dysbiotic infant mice 10 d.p.i. Areas highlighted by broken white lines represent lung damage. **G)** lung injury scores as per American Thoracic Society Guidelines from either control or dysbiotic infant mice 10 d.p.i. [Solid lines, mean; dotted lines, quartiles, * indicated *p*-values < 0.05, Student’s t-test. Solid lines, mean; dotted lines, quartiles]. **H)** Kaplan-Meier plot of the fraction of control and dysbiotic infant mice meeting the euthanasia criteria at indicated times post-infection. [n=8 in each experimental group, * p-value < 0.05, Mantel-Cox log-rank test].

**Fig. S3:**
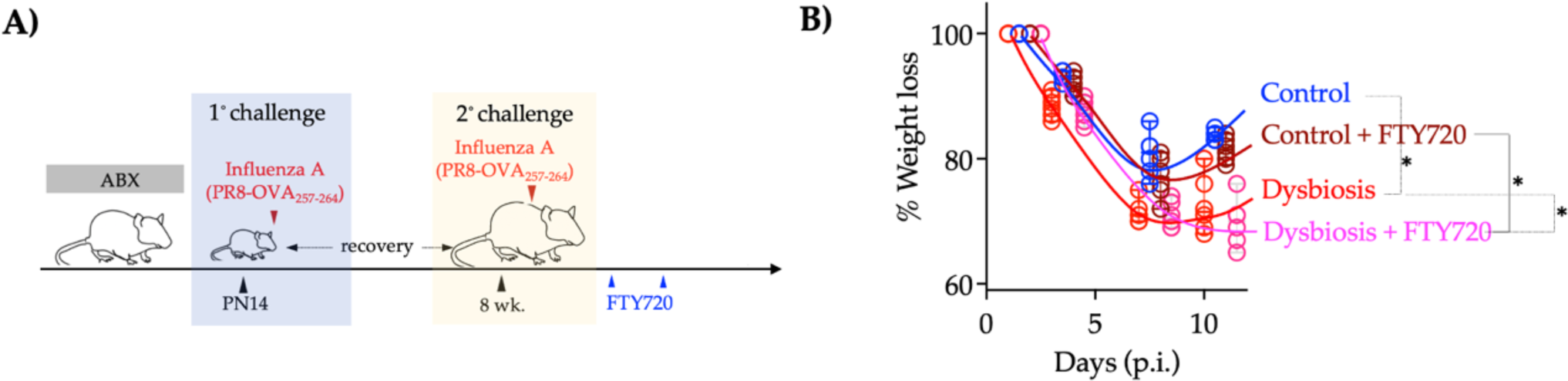
Dysbiotic infant mice mount less robust lung-localized CD8^+^ T cell responses to influenza A (related to Fig. 1). **A)** Experimental approach. Pregnant C57/B6 dams were treated with a cocktail of antimicrobials from embryonic day (E) 15 to postnatal day (PN) (dysbiosis) or with saline (control). Infant mice in each experimental group were challenged with a sublethal dose of murine-adapted influenza A H1N1 strain-PR8, expressing OVA_257–264_ epitope (PR8-OVA) [10^2^ TCID50] via the intranasal (i.n.) route on PN14. Infant mice were allowed to recover for six weeks and challenged with PR8-OVA via i.n. route. Some mice were treated with FTY720 daily throughout the infection. **B)** Weight change at the indicated time after challenge with PR8-OVA [n=5, * *p*-values < 0.05, one-way ANOVA with Tukey’s correction for multiple comparisons. Solid lines, mean; dotted lines, quartiles).

**Fig. S4:**
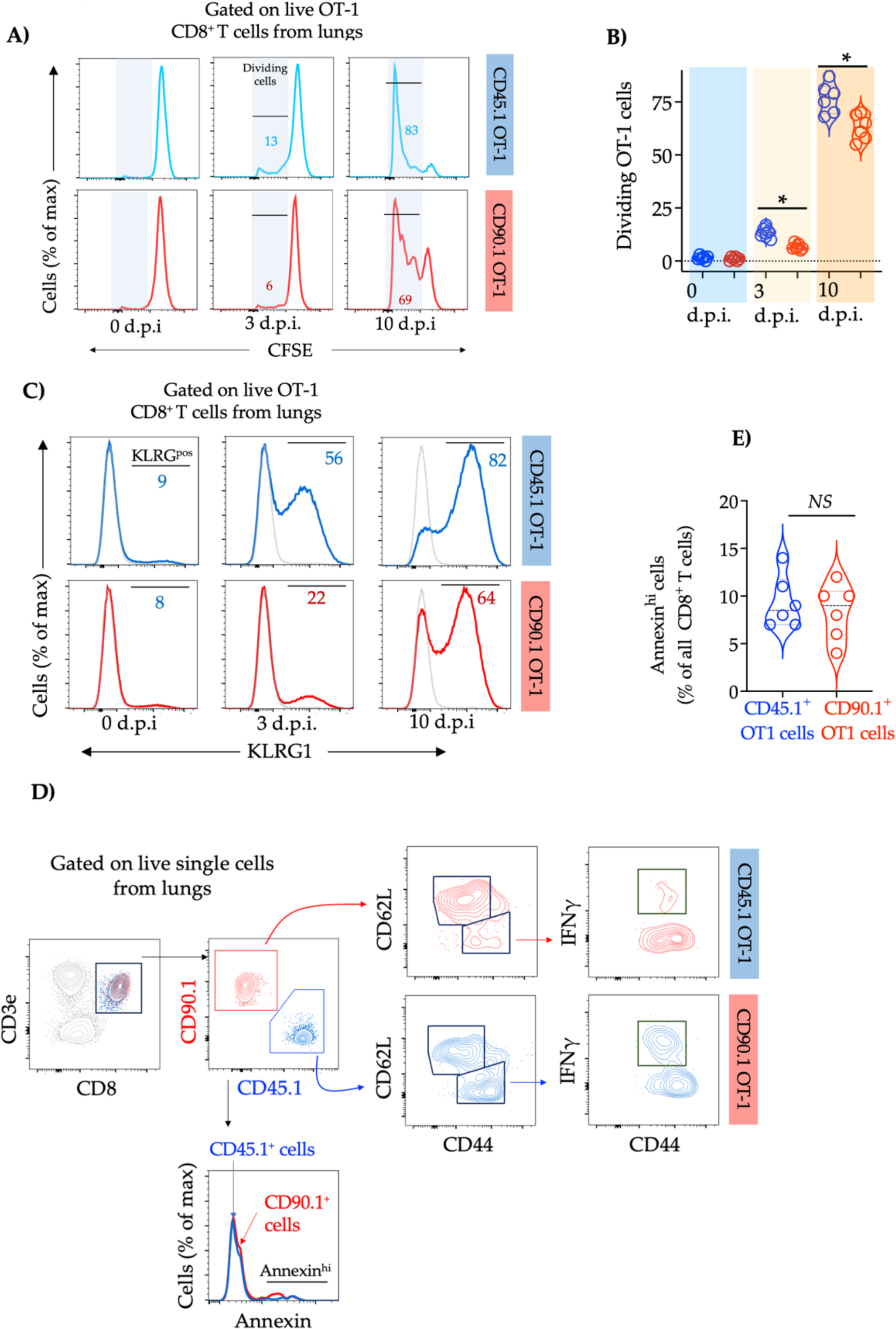
Cell-intrinsic defects in the generation of lung-localized effector T cells in dysbiotic infant mice. (Related to Fig. 1). **A)** Pregnant OT-1 transgenic dams, on either CD90.1 or CD45.1 background, were treated with a cocktail of antimicrobials as described in Fig. 1A. Equal numbers of OT-1 CD8^+^ T cells from control (CD45.1) or dysbiotic (CD90.1) donors (PN13) were labeled with CFSE and adoptively co-transferred into age-matched recipient mice. Twelve hours later, the recipient mice were challenged with influenza [PR8-OVA, 10^2^ TCID50]. Representative histogram of CFSE expression in control (CD45.1) or dysbiotic (CD90.1) OT-1 cells in individual recipient mice at indicated times p.i. **B**) proportion of divided control (CD45.1) or dysbiotic (CD90.1) OT-1 cells in individual recipient mice at indicated times p.i. [Solid lines, mean; dotted lines, quartiles, n=7, * indicated *p*-values < 0.05, Student’s t-test]. **C**) Representative histogram of KLRG1 expression in OT1 CD90.1 (dysbiotic) or OT-1 45.1 (control) CD8^+^ T cells. Proportion of KLRG1^+^ CD8^+^ T cells is indicated. **D**) Flow cytometry analysis of CD44^hi^CD62L^lo^INFγ ^+^ cells in OT1 CD90.1 (dysbiotic) and OT-1 45.1 (control) CD8^+^ T cells. Representative bi-axial plots. **E)** The proportion of Annexin high cells in control (OT-1 CD45.1) and dysbiotic (OT-1 CD90.1) CD8^+^ T cells [n=6, ns, paired student’s t-test. Solid lines, mean; dotted lines, quartiles].

**Fig. S5:**
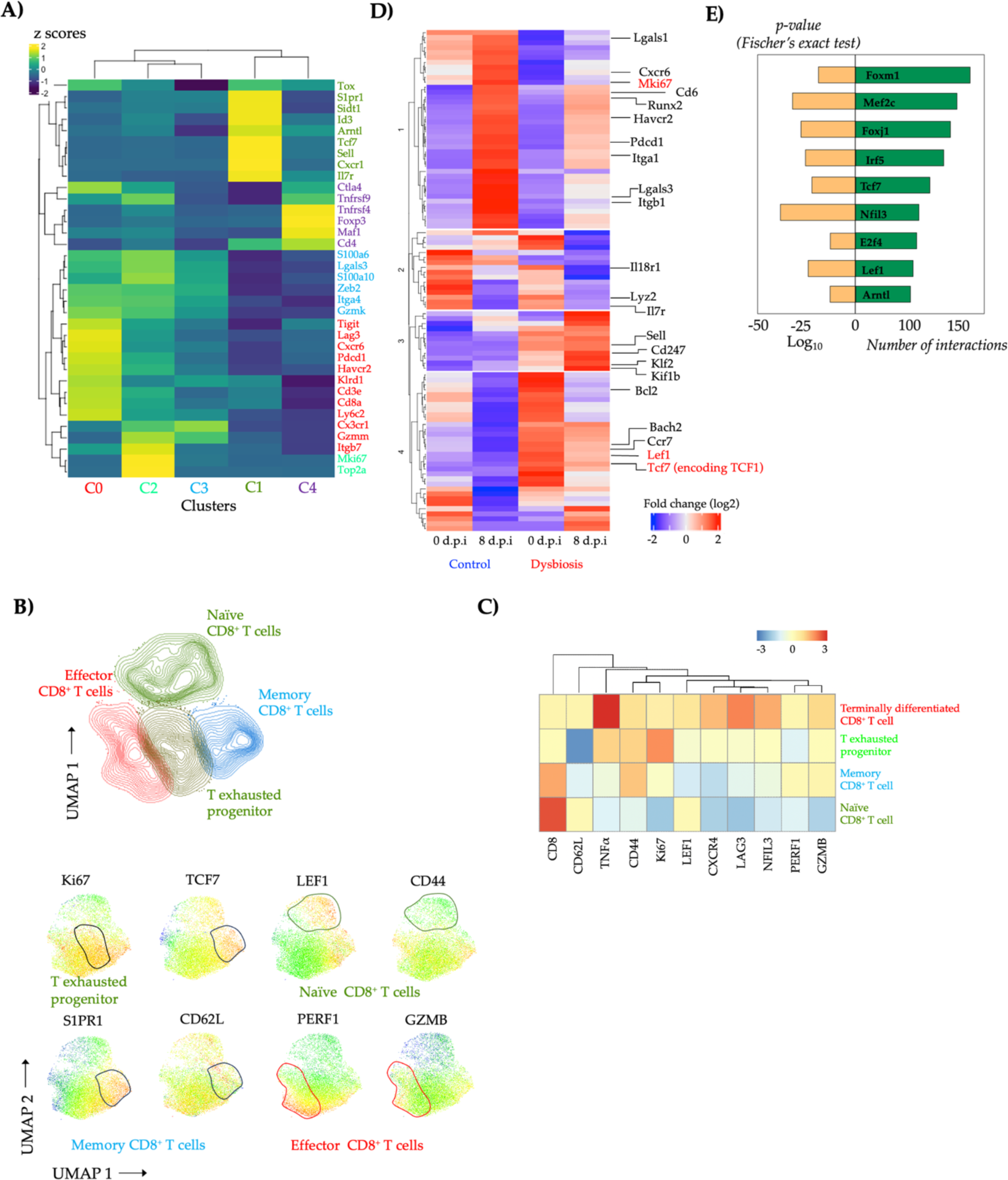
Altered regulatory programs in lung-localized CD8^+^ T cells in dysbiotic infant mice (Related to Fig. 2). **A**) Lung from control and dysbiotic infant mice (n=4, 2 in each group) was obtained at 7 days post-infection (d.p.i). Lung samples were dissociated into cell suspensions, magnetically enriched (EPCAM^-^CD31^-^CD45^+^CD8^+^), and used for single-cell RNA sequencing (scRNAseq). Row-scaled expression of genes uniquely expressed in each cluster. k-means clustering was used to arrange subjects and transcripts. **B**) Lung samples dissociated into cell suspensions were used for flow cytometry analysis. Unsupervised analysis of live, single live CD8 T^+^ cells using a self-organizing map (SOM). CD8^+^ T cell clusters identified by SOM were mapped to uniform manifold approximation and projection (UMAP) embedding and colored by naïve, effector, memory, and exhausted progenitor clusters. UMAP projection showing expression of key phenotypic markers. **C**) Row-scaled mean fluorescent intensity (MFI) of key phenotypic markers in the indicated cluster. k-means clustering of phenotypic markers [Benjamini and Hochberg-adjusted *p*-values < 0.01, log_2_ fold change > 2, Wald’s test]. **D**) Row-scaled expression of the highest differentially expressed genes (DEG) in control or dysbiotic infant mice. k-means clustering was used to arrange subjects and transcripts. **E**) Transcriptional activity score [negative -log two-sided *p*-value of TF enrichment per cell type by Fisher exact test] for TFs. Yellow bar = TAS, and green bar = # of regulated genes.

**Fig. S6:**
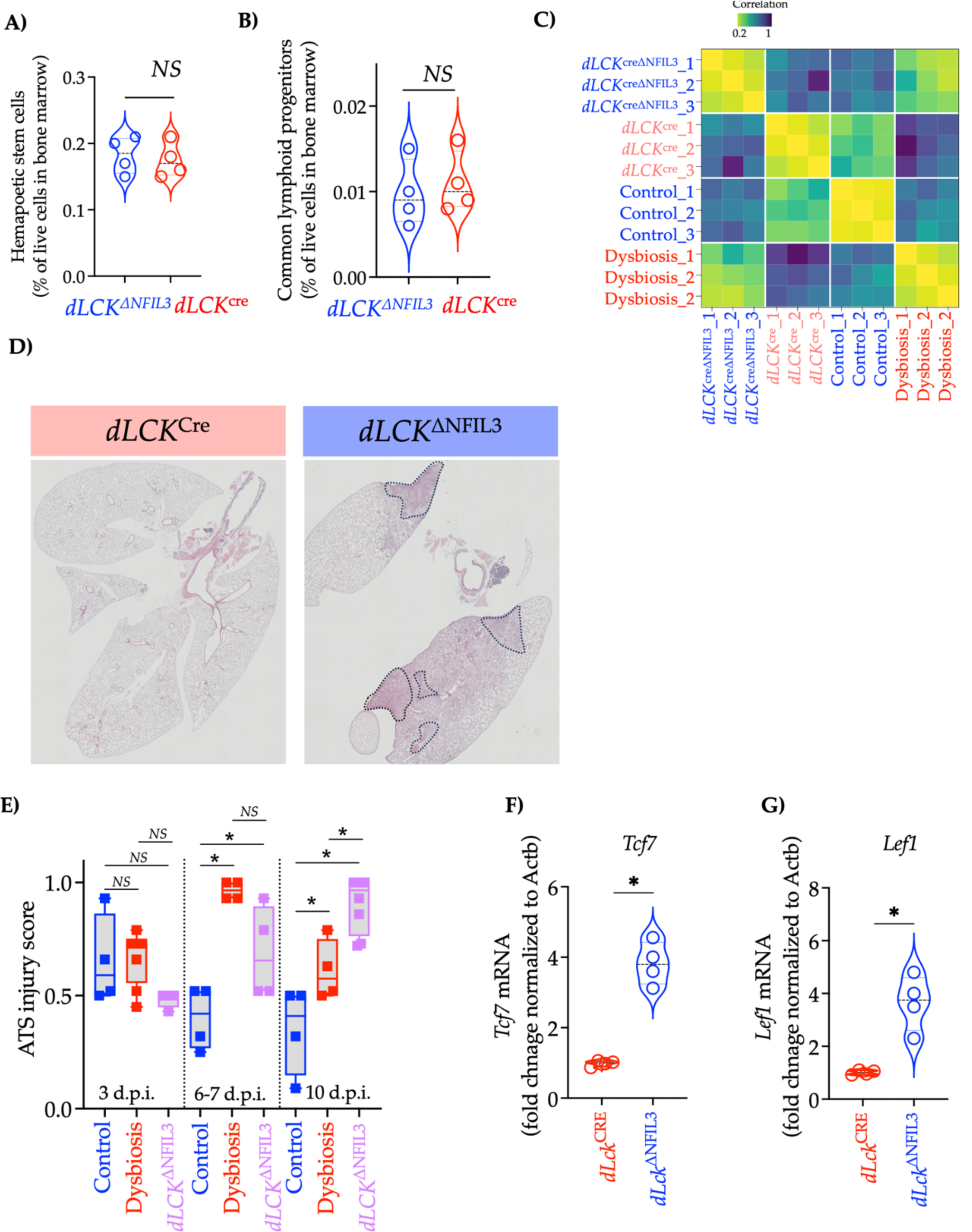
T cell-specific deletion of NFIL3 remodels the lung-localized CD8^+^ T cell compartment. (Related to Fig. 4) **A)** Frequencies of HSC (identified as Lin^-^c-Kit^+^ cells) and **B**) CLP (identified as Lin-Flt3^+^c-Kit^mid^Sca-1^mid−^CD48^+^ cells) in the bone marrow of *dLck*^ΔNfil3^ or *dLck*^cre^ infant mice (PN14). [n =6, NS – not significant, paired student’s t-test, violin plots, Solid lines, mean; dotted lines, quartiles]. **C**) Pairwise Euclidean distances between CD8^+^ T cells in the lungs of dysbiotic, *dLck*^ΔNfil3^, and *dLck*^cre^ infants. **D**) Photomicrographs of hematoxylin and eosin (H&E) stained sections of lungs from either *dLck*^ΔNfil3^ or *dLck*^cre^ infant mice 10 d.p.i. Areas highlighted by broken white lines represent lung damage. **E**) Lung injury scores as per American Thoracic Society Guidelines from *dLck*^ΔNfil3^ or control or dysbiotic infant mice at indicated times post-infection. Data for control infant mice was re-used from Fig. S3D. [Solid lines, mean; dotted lines, quartiles, * indicated *p*-values < 0.05, one-way ANOVA with Tukey’s correction for multiple comparisons. Solid lines, mean; dotted lines, quartiles]. **F**) Fold change in *Tcf7* or **G**) *Lef1* mRNA in lung CD8^+^ T cells from *dLck*^ΔNfil3^ and *dLck*^cre^ infant mice (PN14) [n=3, * indicated *p*-values < 0.05, Student’s t-test. Solid lines, mean; dotted lines, quartiles].

**Fig. S7:**
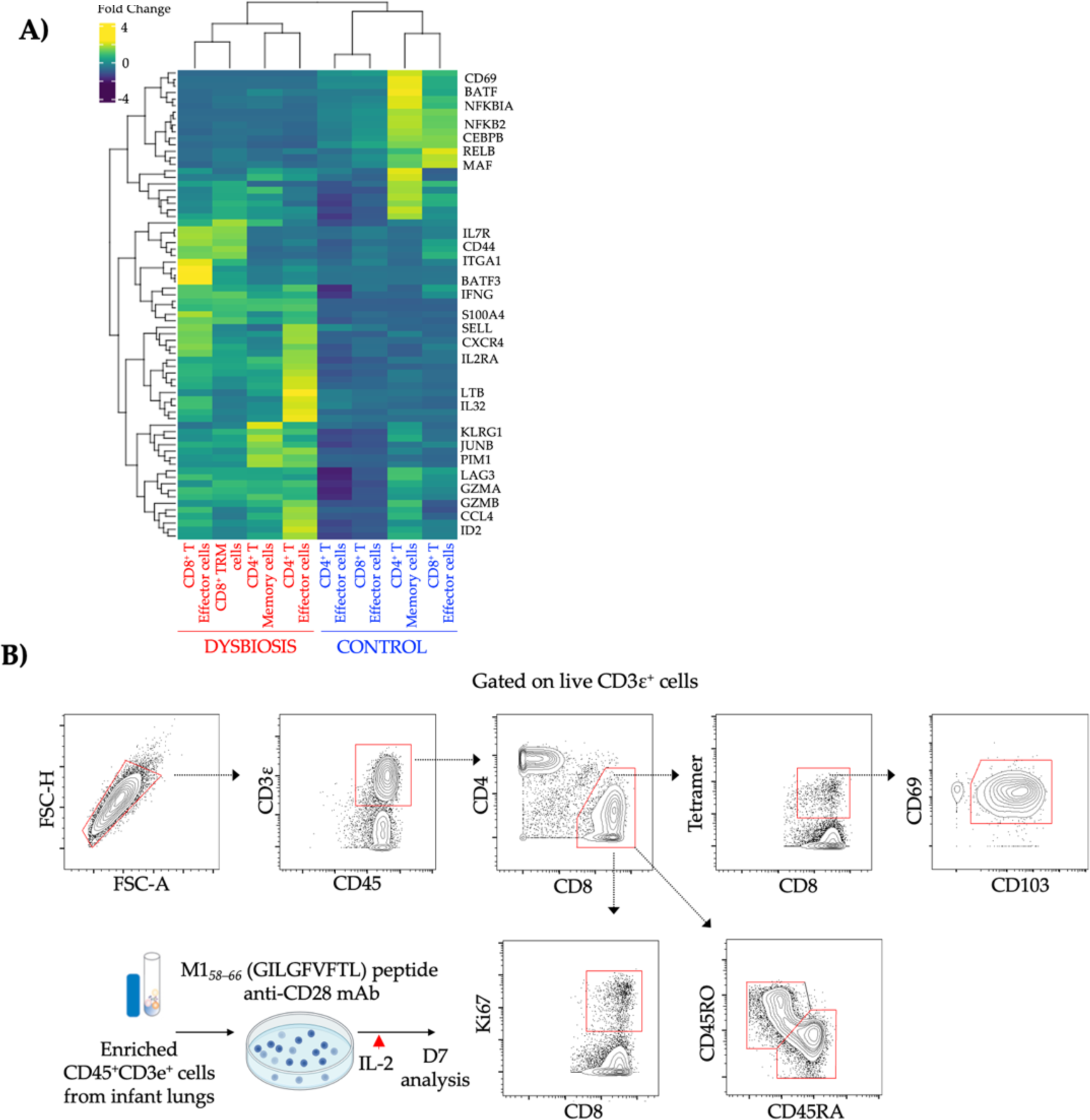
Cell-intrinsic defects in the generation of lung-localized effector T cells in dysbiotic human infants (Related to Fig. 3). **A)** Enriched (CD45^+^CD3ε ^+^) single-cell suspension from control (no-ABX exposed) or dysbiotic (ABX-exposed) human infants were used for single-cell RNA sequencing (scRNAseq). Row-scaled expression of genes uniquely expressed in each cluster. k-means clustering was used to arrange subjects and transcripts. **B**) Enriched (CD45^+^CD3ε ^+^) single-cell suspension from control and dysbiotic human infants were co-incubated with >M158>–66 (GILGFVFTL) peptide and anti-CD28 mAb, followed by stimulation with IL-2 and analyzed by flow cytometry after seven days. Gating strategy related to Fig. 5H. Representative bi-axial plots.

**Fig. S8:**
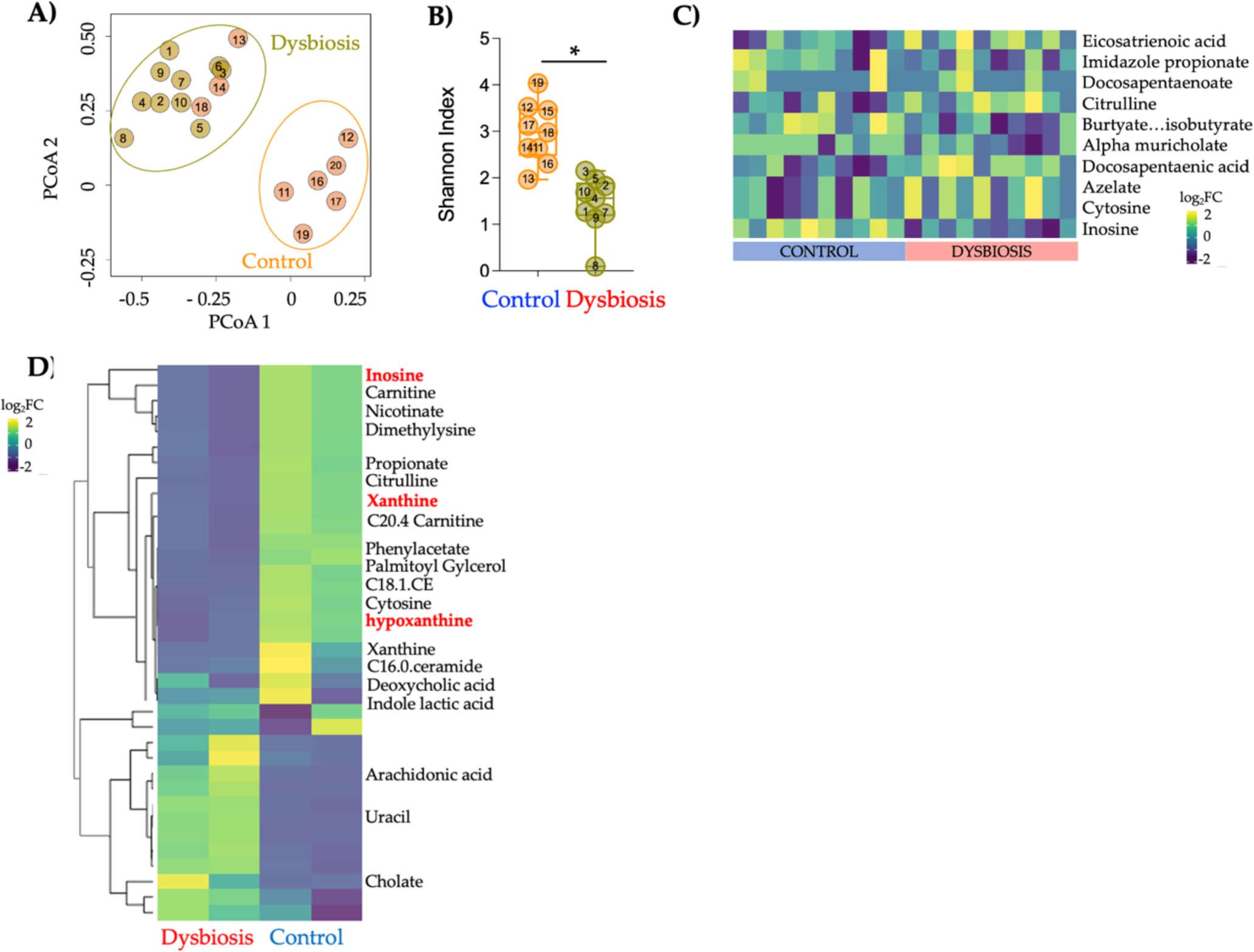
Microbiome-derived inosine mediates CD8^+^ T cell proliferation, effector differentiation, and host resistance to influenza (Related to Fig. 5). **A)** Principal Component (PCA) of fecal microbial communities of dysbiotic and control human infants, colored by experimental group. **D**) β-diversity (unweighted UniFrac) of fecal microbiota of dysbiotic and control human infants [n=10, * *p*-values < 0.05, Tukey bars, lines at mean and quartiles]. **C)** Row-scaled abundance of metabolites in intestinal contents predicted based on metabolic pathways identified by shotgun metagenomic sequencing between dysbiotic and control human infants. **D**) Row-scaled abundance of metabolites differentially abundant expressed in serum of dysbiotic and control infant mice. k-means clustering was used to arrange metabolites (Bonferroni-adjusted *p*-value < 0.05).

**Fig. S9:**
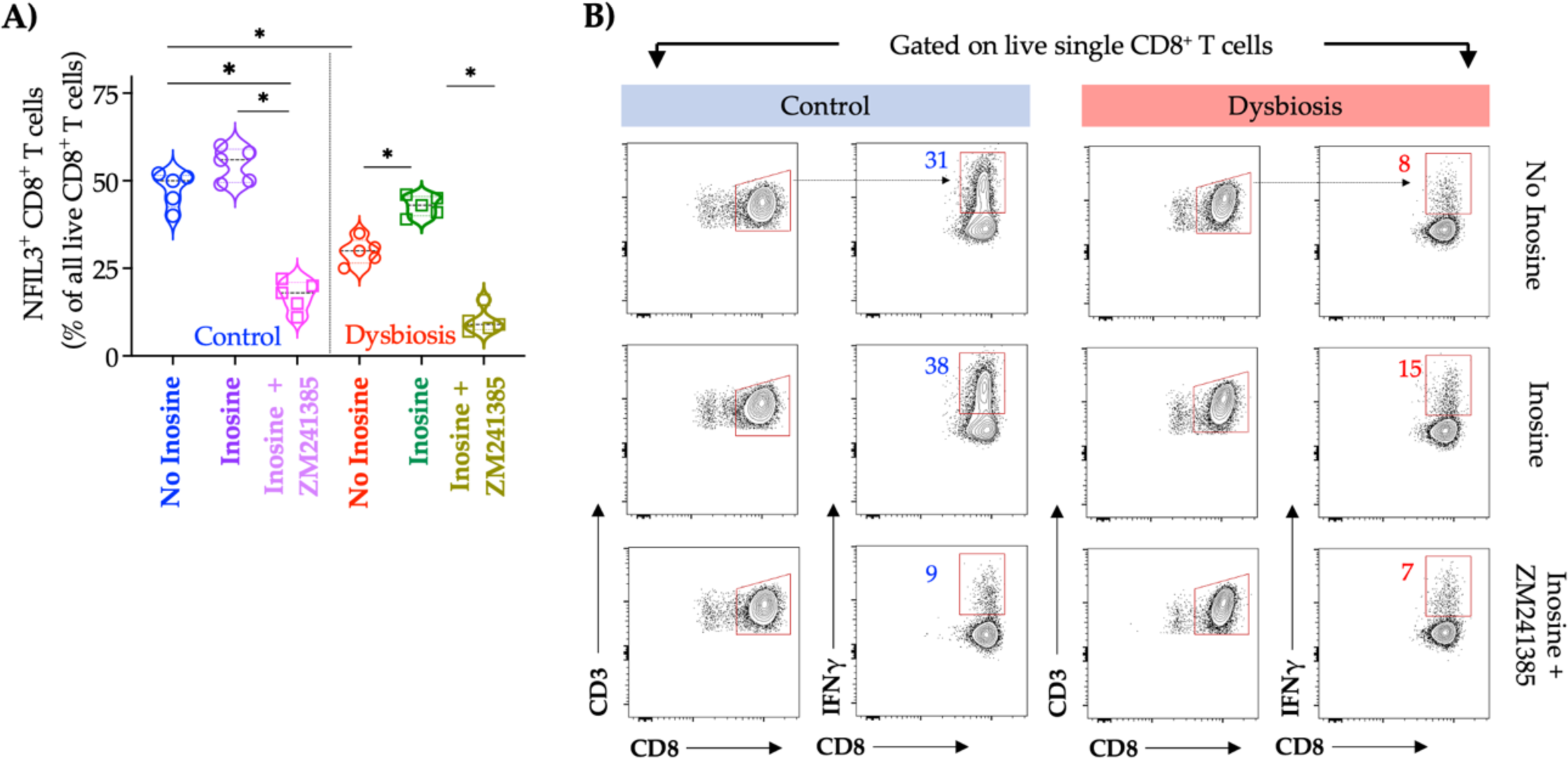
Microbiome-derived inosine mediates CD8^+^ T cell proliferation, effector differentiation, and host resistance to influenza (Related to Fig. 5). **A)** Enriched (CD3ε^+^ CD8^+^) single-cell suspension from control and dysbiotic infant mice were co-incubated with inosine with or without an A2AR antagonist (ZM241385). Proportion of NFIL3^+^ CD8^+^ T cells in indicated experimental groups [n=5, * indicated *p*-values < 0.05, one-way ANOVA with Tukey’s correction for multiple comparisons. Solid lines, mean; dotted lines, quartiles]. **B)** Representative biaxial plots of IFNγ in CD8^+^ T cells in each experimental group. The proportion of IFNγ^+^ CD8^+^ T cells is indicated.

**Fig. S10:**
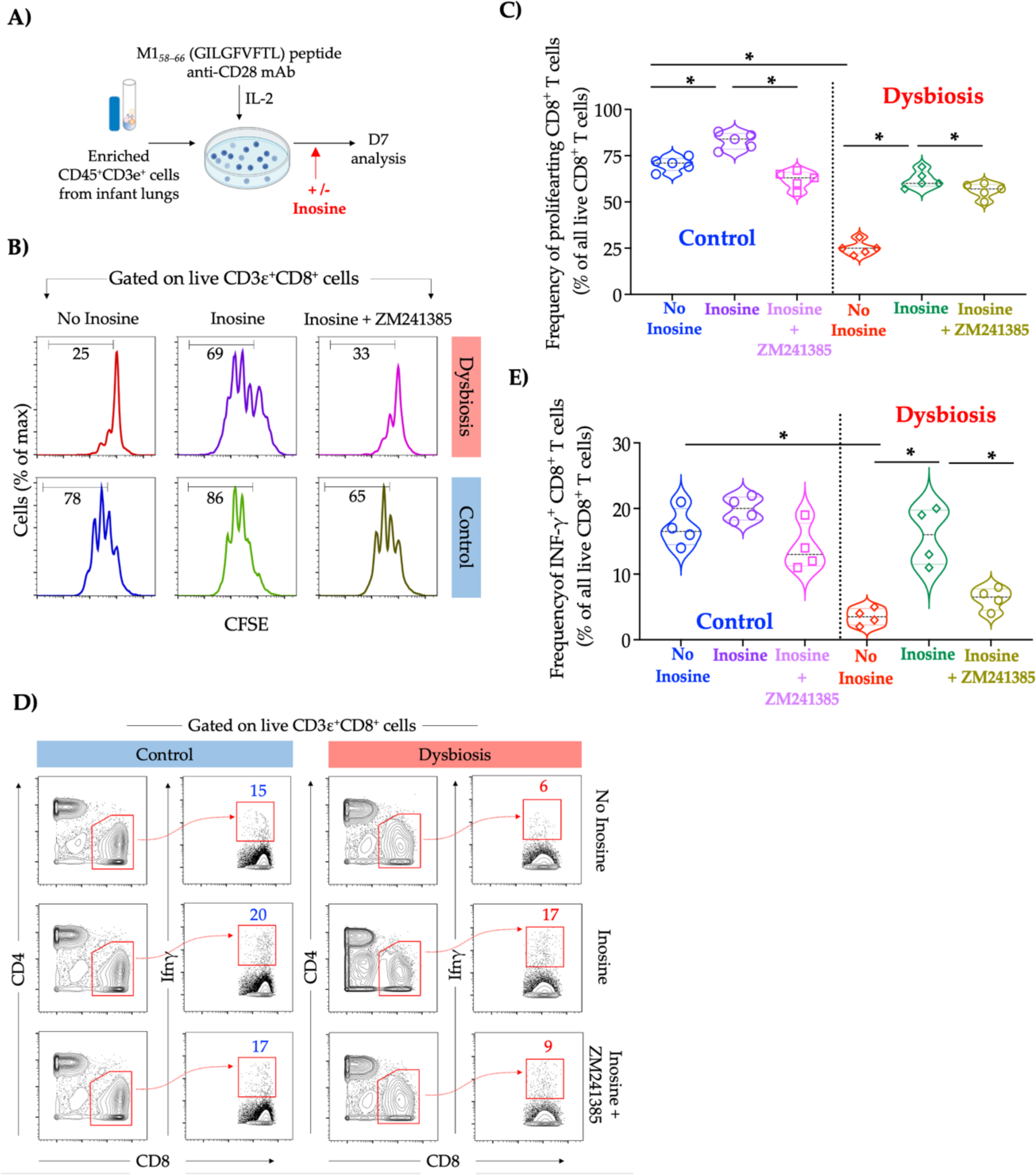
Microbiome-derived inosine is essential for robust lung-localized anti-viral response and NFIL3-directed remodeling of the lung-T cell compartment in dysbiotic human infants (Related to Fig. 5). **B)** Enriched (CD45^+^CD3ε ^+^) single-cell suspension from control and dysbiotic human infants were co-incubated with >M158>–66 (GILGFVFTL) peptide and anti-CD28 mAb, followed by stimulation with IL-2 in the presence or absence of inosine and analyzed by flow cytometry after seven days. **C)** Representative histogram of CFSE expression in CD8^+^ T cells in each experimental group. The proportion of proliferating CD8^+^ T cells is indicated. **C)** Proportion of proliferating CD8^+^ T cells in each experimental group [n=5, * indicated *p*-values < 0.05, one-way ANOVA with Tukey’s correction for multiple comparisons. Solid lines, mean; dotted lines, quartiles]. **D)** Representative biaxial plots of IFNγ in CD8^+^ T cells in each experimental group. The proportion of IFNγ^+^ CD8^+^ T cells is indicated. **E)** Proportion of IFNγ^+^ CD8^+^ T cells in each experimental group [n=5, * indicated *p*-values < 0.05, one-way ANOVA with Tukey’s correction for multiple comparisons. Solid lines, mean; dotted lines, quartiles].

**Fig. S11:**
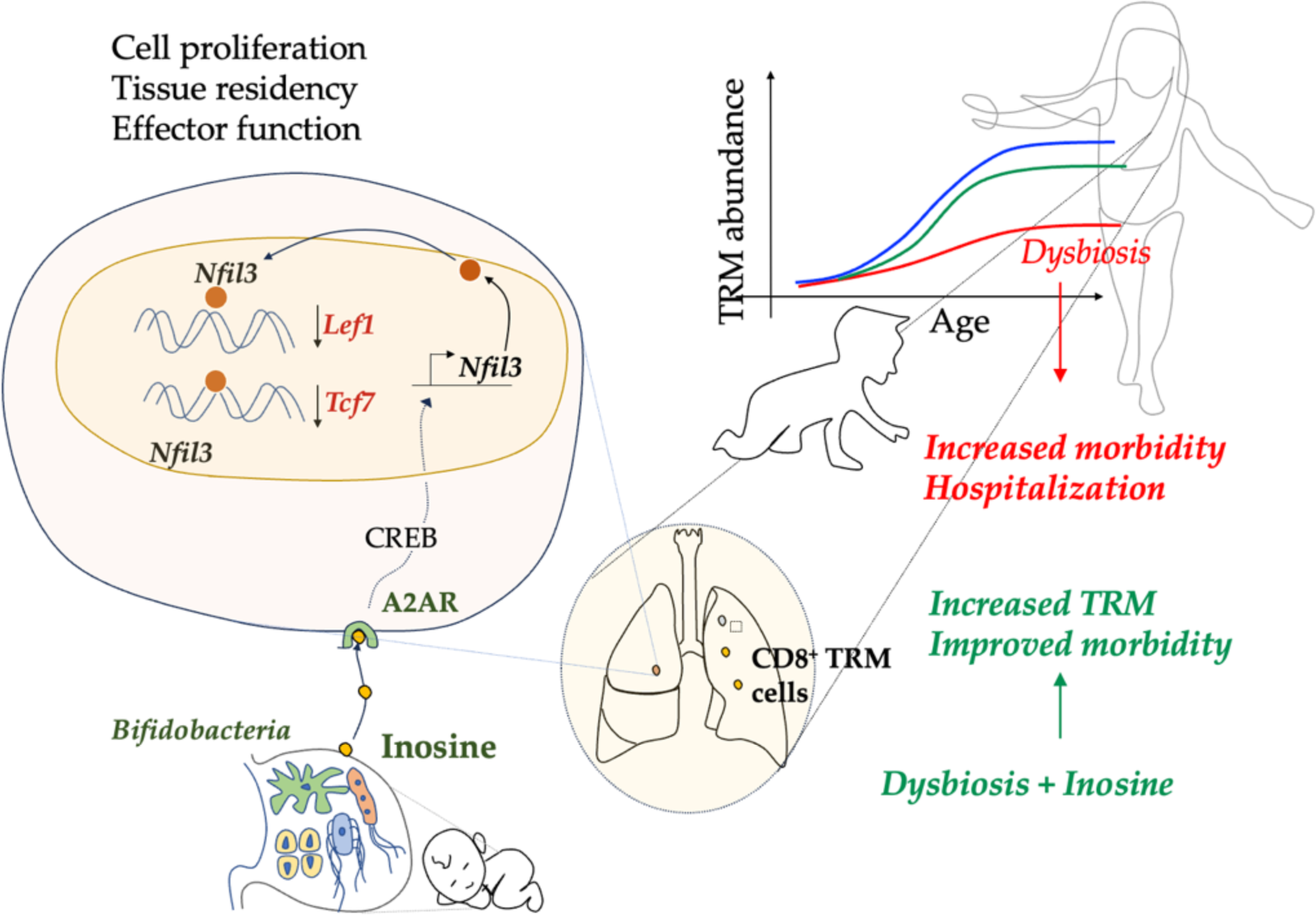
Microbiota-derived inosine programs protective CD8^+^ T cell responses against influenza in newborns. Influenza virus susceptibility in dysbiotic infant mice is caused by CD8^+^ T cell hyporesponsiveness and diminished persistence as tissue-resident memory cells. Nuclear factor interleukin 3 (NFIL3) repressed memory differentiation of CD8^+^ T cells in dysbiotic mice by epigenetic regulation of T cell factor 1 (TCF 1) expression. Pulmonary CD8^+^ T cells from dysbiotic human infants share these transcriptional signatures and functional phenotypes. Mechanistically, intestinal inosine was reduced in dysbiotic human infants and newborn mice, and inosine replacement reversed epigenetic dysregulation of *Tcf7* and increased memory differentiation and responsiveness of pulmonary CD8^+^ T cells.

**Table S1:** Demographic characteristics of human infants involved in single cell RNA sequencing and analysis (scRNA seq), spectral cytometry, and ex vivo assays.

| **Age** | **Diet** | **Race** | **Group** | **Ethnicity** | **Sex** | **Lung Samples** | **Antibiotic courses (>5 days)** | **RTI history** |
| --- | --- | --- | --- | --- | --- | --- | --- | --- |
| 5-6 months | Mixed | White | Dysbiosis | Hispanic or Latino | M | YES | 4 | YES |
| 7-8 months | Mixed | White | Dysbiosis | Not Hispanic or Latino | F | YES | 5 | YES |
| 3-4 months | Mixed | White | Dysbiosis | Hispanic or Latino | F | YES | 3 | YES |
| 4-5 months | Mixed | White | Dysbiosis | Not Hispanic or Latino | F | YES | 4 | YES |
| 4-5 months | Breast Milk | White | Control | Not Hispanic or Latino | F | YES | 0 | YES |
| 5-6 months | Mixed | Black African American | Control | Not Hispanic or Latino | M | YES | 1 | YES |
| Less than 1 month | Infant Formula | Black African American | Control | Not Hispanic or Latino | F | YES | 0 | No |
| 5-6 months | Mixed | White | Control | Hispanic or Latino | F | YES | 0 | YES |

**Table S2:** Demographic characteristics of human infants involved in microbiome studies.

| Age | Diet | Race | Group | Ethnicity | Sex |
| --- | --- | --- | --- | --- | --- |
| 1-2 months | Mixed | Black African American | Dysbiosis | Not Hispanic or Latino | Male |
| 5-6 months | Mixed | White | Dysbiosis | Hispanic or Latino | Male |
| 7-8 months | Mixed | White | Dysbiosis | Not Hispanic or Latino | Female |
| Less than 1 month | Breast Milk | White | Dysbiosis | Hispanic or Latino | Female |
| 3-4 months | Mixed | White | Dysbiosis | Hispanic or Latino | Female |
| 1-2 months | Infant Formula | White | Dysbiosis | Not Hispanic or Latino | Female |
| 2-3 months | Breast Milk | White | Dysbiosis | Not Hispanic or Latino | Male |
| 4-5 months | Mixed | White | Dysbiosis | Not Hispanic or Latino | Female |
| 1-2 months | Breast Milk | White | Dysbiosis | Not Hispanic or Latino | Female |
| Less than 1 month | Breast Milk | White | Dysbiosis | Not Hispanic or Latino | Male |
| 2-3 months | Breast Milk | White | Control | Not Hispanic or Latino | Male |
| 3-4 months | Breast Milk | White | Control | Hispanic or Latino | Male |
| 4-5 months | Breast Milk | White | Control | Not Hispanic or Latino | Female |
| 5-6 months | Mixed | Black African American | Control | Not Hispanic or Latino | Male |
| Less than 1 month | Breast Milk | White | Control | Not Hispanic or Latino | Male |
| Less than 1 month | Infant Formula | Black African American | Control | Not Hispanic or Latino | Female |
| Less than 1 month | Breast Milk | Black African American | Control | Not Hispanic or Latino | Male |
| Less than 1 month | Breast Milk | Black of African American | Control | Not Hispanic or Latino | Male |
| 5-6 months | Mixed | White | Control | Hispanic or Latino | Female |
| Less than 1 month | Breast Milk | White | Control | Not Hispanic or Latino | Female |

**Table S3:**
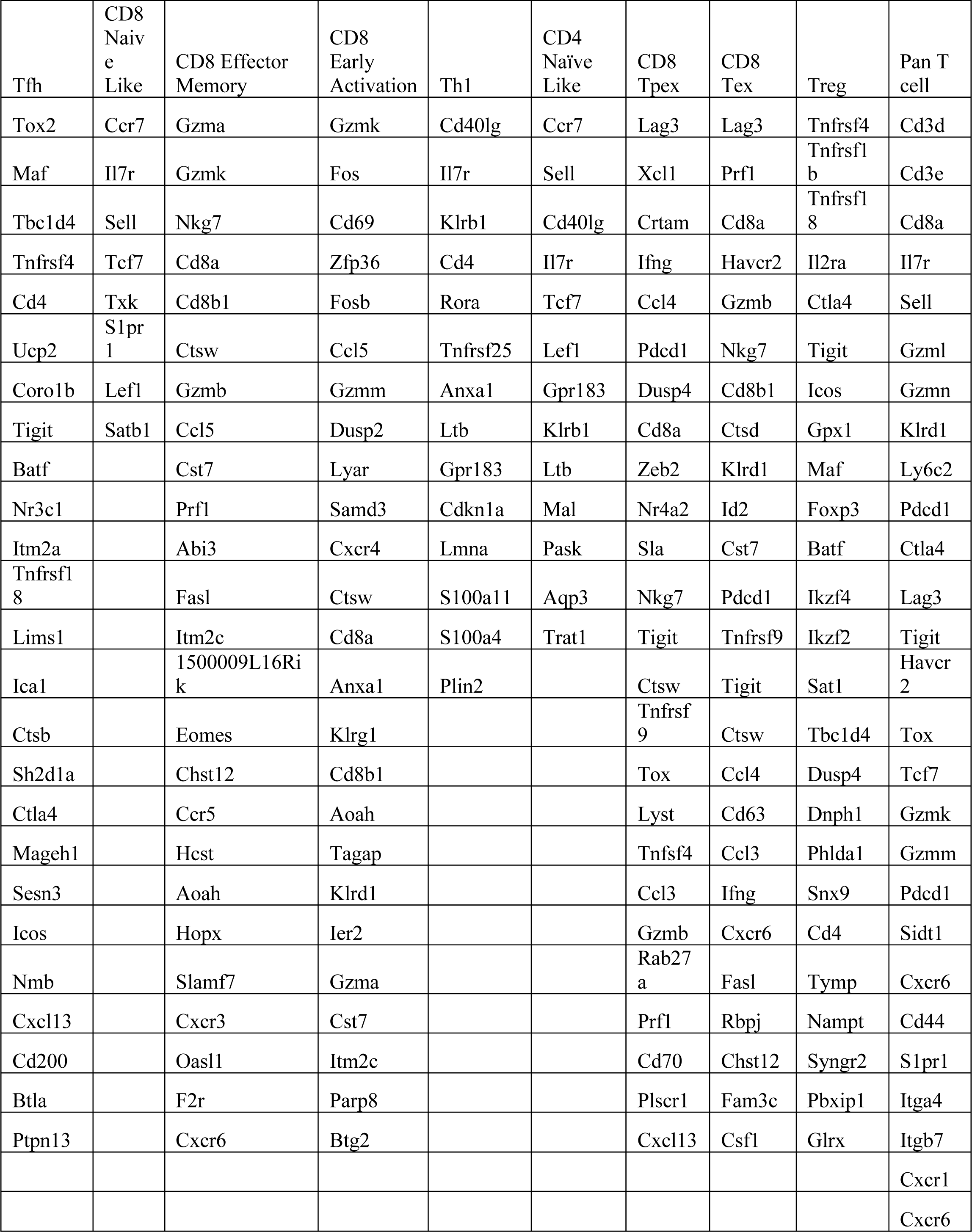
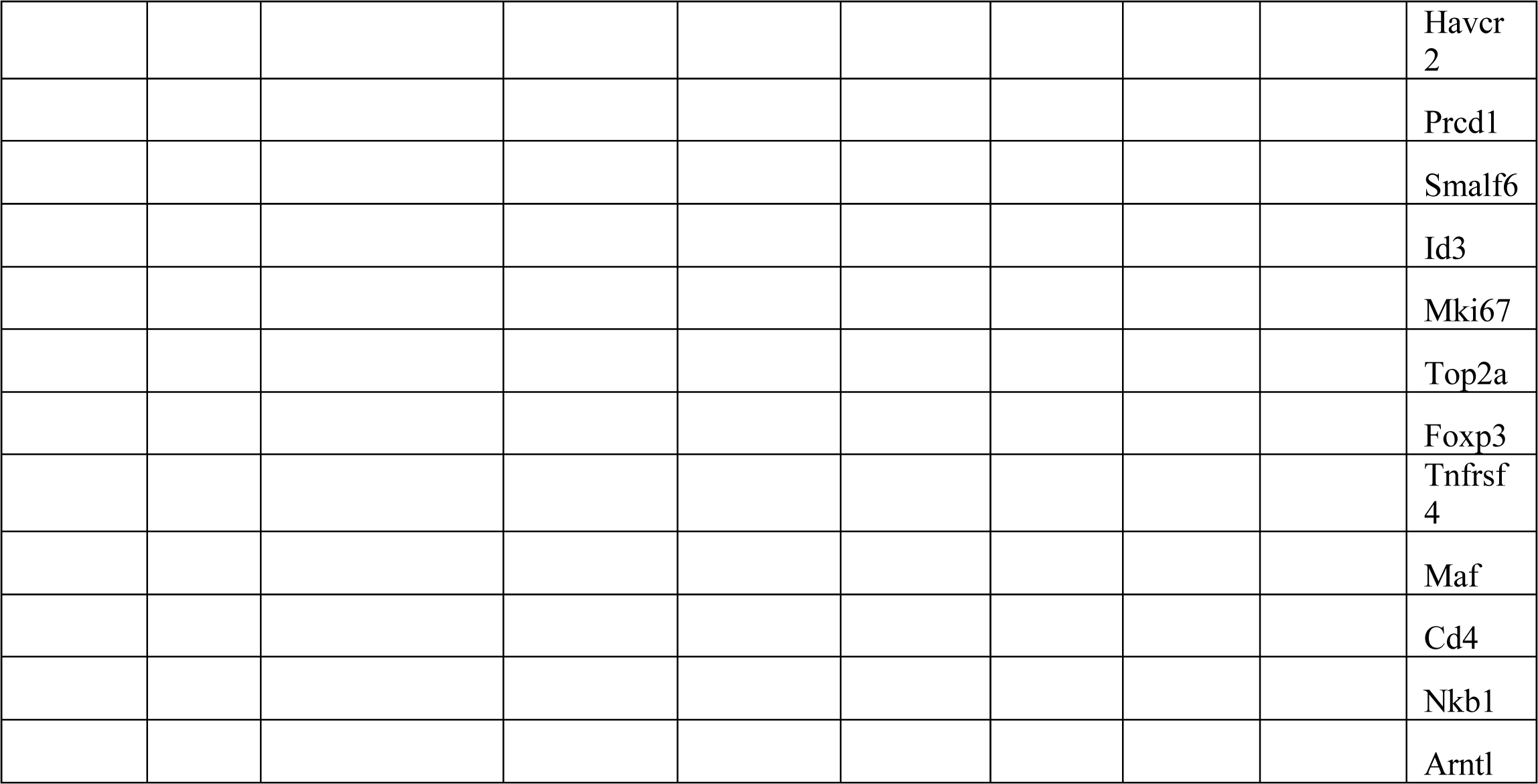
Cell state-specific gene signatures.

**Table S4:**
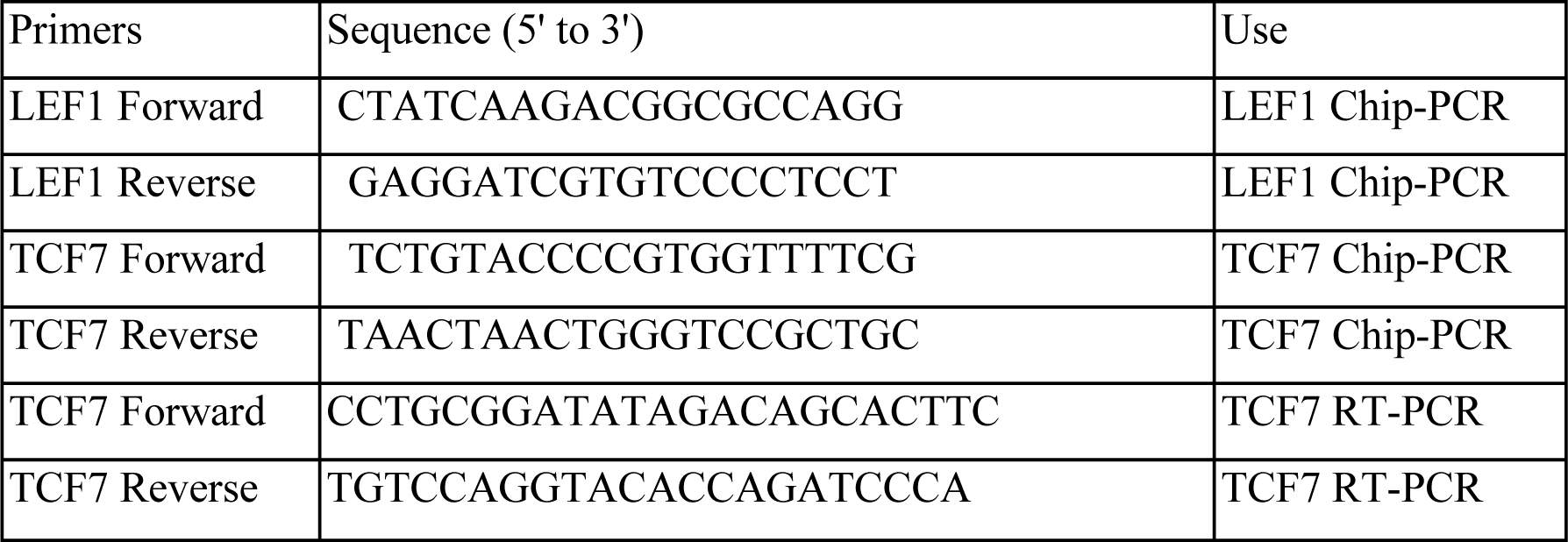
Primers used for Chip-PCR and RT PCR.

**Table S5:** Antibodies used for flow cytometry studies in mice.

| Target | Clone | Company | Cat/Ref# | Dilution used |
| --- | --- | --- | --- | --- |
| Galectin 3 | M3/38 | Novus Bio | NBP1-43313AF350 | 1:100 |
| CXCR4 (CD184) | 2B11/CXCR4 | BD Bioscience | 561735 | 1:100 |
| Galectin 1 | RM0081-9J26 | Novus Bio | NBP1-21721AF350 | 1:100 |
| NFIL3 | S2M-E19 | eBioscience | 12-5927-80 | 1:500 |
| CD45 | 30-F11 | BioLegend | 103128 | 1:100 |
| CD8a | 53-6.7 | BioLegend | 100751 | 1:100 |
| IFN-g | XMG1.2 | BD Bioscience | 566366 | 1:100 |
| TNF | MP6-XT22 | BD Bioscience | 566511 | 1:100 |
| CD62L | MEL-14 | BioLegend | 104445 | 1:100 |
| CD44 | IM7 | BioLegend | 103039 | 1:100 |
| S1P1 | 713412 | Novus Bio | FAB7089C | 1:100 |
| CD4 | GK1.5 | Novus Bio | NBP2-25191AF350 | 1:100 |
| p-CREB | 87G3 | Cell Signaling Technology | 9187 | 1:50 |
| LEF1 | C12A5 | Cell Signaling Technology | 14022 | 1:100 |
| Ki-67 | 16A8 | BioLegend | 652413 | 1:100 |
| Perforin | S16009A | BioLegend | 154315 | 1:100 |
| Granzyme B | NGZB | eBioscience | 25-8898-82 | 1:100 |
| CD69 | H1.2F3 | BioLegend | 104502 | 1:100 |
| CD103 | W19396D | BioLegend | 110902 | 1:100 |
| CD3e | 145-2C11 | BioLegend | 100303 | 1:100 |
| CD45.1 | A20 | eBioscience | 47-4321-82 | 1:100 |
| CD45.2 | 104 | BioLegend | 109821 | 1:100 |
| CD90.1 | S20007C | BioLegend | 109002 | 1:100 |
| KLRG1 | 2F1/KLRG1 | BioLegend | 138405 | 1:100 |
| Sca-1 | D7 | BioLegend | 108111 | 1:100 |
| c-kit | ACK2 | BioLegend | 135107 | 1:100 |
| FLT3 | A2F10 | BioLegend | 135305 | 1:100 |
| CD48 | HM48-1 | BioLegend | 103409 | 1:100 |
| TCF1 | C63D9 | Cell Signaling Technology | 9066S | 1:100 |
| LAG3 | Polyclonal | Novus Bio | AF3328 | 1:100 |
| SIINFEKL tetramer | MKb-001 | Tetramer Shop (discontinued) | 20220121F | 1:100 |

## Supplementary Materials for

### Materials and Methods

#### Experimental Model and Subject Details

**Mice:** The overall goal of this study was to test if disruption to the early-life microbiota would disrupt pulmonary T cell responses and cause worsened outcomes to viral respiratory pathogens. The Institutional Animal Care and Use Committees (IACUCs) at Cincinnati Children’s Hospital Medical Center (CCHMC) approved all the animal studies (IACUC protocol number 2023-0053), which were carried out by the Association for the Assessment and Accreditation of Laboratory Animal Care (AAALAC)-accredited specific pathogen-free (SPF) animal facility at CCHMC. We obtained *dLCK*^cre^ (#12837)(*1*), OT-1 (#003831)(*2, 3*), C57B/6J (#000664), B6.PL-*Thy1^a^*/CyJ (expressing CD90.1) (#000406)(*4*), and B6.SJL-*Ptprc^a^ Pepc^b^*/BoyJ (expressing CD45.1) (#002104)(*5*) from Jackson Laboratory. *Nfil3*^flox^ mice were a gift from Drs. Masato Kubo (*6*). We bred *dLCK*^cre^ and *Nfil3*^flox^ to generate *dLCK^ΔNFIL3^* mice. OT-1 mice were backcrossed with either B6.PL-*Thy1^a^*/CyJ or B6.SJL-*Ptprc^a^ Pepc^b^*/BoyJ mice to generate OT-1 mice expressing CD45.1 or CD90.1 respectively.

#### Human Studies

The Institutional Review Board (IRB) at Cincinnati Children’s Hospital Medical Center (CCHMC) approved all human subject studies (IRB protocol numbers, 2018-0852 and 2017-3726). The study does not qualify as human subjects research as determined by CCHMC IRB because tissue samples were obtained from deceased individuals. Demographics of human subjects are listed in **Table S1 and S2**.

#### Antibiotic exposure

Pregnant female mice were randomized to receive either sucralose drinking water mixed with equal weights by volume of ampicillin, gentamicin, and vancomycin (1 mg/mL) or sucralose drinking water alone starting from embryonic day 15 until day 5 of life for the newborn. After birth, neonatal mice from multiple litters were pooled and redistributed to control for the founder effect and to minimize in-cage variations. Newborn mice and their dams were exposed to a standard light cycle (lights on for 12 h starting at 08:00). We used mice between ages P14 and eight weeks of age and appropriate age-, sex- and genetic strain-matched controls to account for any variations in data. Published data in newborn mice was used to estimate that seven animals in each group would be sufficient to detect a 20% difference in morbidity with 80% power and an α of 0.05.

#### Experimental studies

Newborn (PN14) or adult (8 wk) mice were anesthetized by inhalation of isoflurane (5% induction/2% maintenance) and infected with weight-adjusted, sub-lethal doses of murine-adapted influenza A H1N1 strain-PR8, expressing OVA_257–264_ epitope (*7*) [referred hereafter as PR8-OVA] (10^2^ TCID50 for infants or 10^3^ TCID50 for adults (heterotypic challange] or X31 (A/Hong Kong/1/68-x31 [H3N2]) (10^4^ TCID50 for adults) via intranasal route. Morbidity after infection was monitored by daily examination and weight assessment. Influenza viral titers in lung homogenates were measured by a 50% tissue culture infective dose (TCID_50_) assay, as previously described(*3*), with titers expressed as the reciprocal of the dilution of lung extract that corresponded to 50% virus growth in Madin–Darby canine kidney cells, as calculated by the Reed– Muench method (*8*).

To delineate the role of circulating vs. lung resident T cells, mice were treated with the sphingosine 1-phosphate receptor-1 agonist FTY720 (25 μg) or sterile saline via drinking water beginning 1 day before infection and continuing daily throughout infection. To interrogate the role of inosine, infant mice were treated with inosine (300 μg g^-1^) on PN10,12 and 14 via intraperitoneal route.

#### Competitive adoptive CD8^+^ T cell co-transfer

We maintained OT-1 mice expressing transgenic T cell receptors recognizing ovalbumin peptide residues 257-264 (OVA_257-264_)(*9*) on either CD90.1 background or CD45.1 background. OT-1 mice on CD90.1 background or CD45.1 background were exposed to perinatal ABX or No ABX, respectively. Pooled lungs from >5 newborn mice (PN14) from either OT-1 mice on a CD90.1 background (ABX exposed) or a CD45.1 background (No ABX) were gently dissociated by mechanical agitation and strained through a 100-μm nylon filter to obtain a single-cell suspension. CD8^+^ T cells were purified by negative magnetic selection with anti-CD8a microbeads (Miltenyi, Cat. no. 130-095-236). This protocol yielded approximately 5×10^5^ CD8^+^ T cells. OT-1 CD8^+^ T cells from control (CD45.1^+^) newborn mice (PN14) or ABX (CD90.1^+^) newborn mice (PN14) were labeled with CFSE per the manufacturer’s protocol (Thermo Fisher Scientific), mixed in the ratio of 1:1, and injected via tail vein injection into C57B6 mice (CD45.2^+^) 1 day before infection. The rate of division and proliferation index was estimated using the precursor method as previously described(*10*).

#### Adoptive transfer of CD8^+^ T cells from *dLCK^ΔNFIL3^* and *dLCK*^cre^ infant mice

Pooled lungs from >5 newborn mice (PN14) from either *dLCK^ΔNFIL3^*or *dLCK*^cre^ infant mice were gently dissociated by mechanical agitation and strained through a 100-μm nylon filter to obtain a single-cell suspension. CD8^+^ T cells were purified by negative magnetic selection with anti-CD8a microbeads. CD8^+^ T cells from either *dLCK^ΔNFIL3^* or *dLCK*^cre^ infant mice labeled with CFSE per the manufacturer’s protocol (Thermo Fisher Scientific) and injected via tail vein into age-matched C57B/6J 1 day before infection. The rate of division and proliferation index was estimated using the precursor method.

#### Tissue processing

Mice were euthanized at either seven- or fourteen-days post-infection. Ten minutes prior to euthanasia, mice were administered 5 µg fluorochrome-conjugated anti-CD45 Ab by intravenous route. Mice were anesthetized with ketamine/xylazine, and 1 ml of PBS was injected into the right ventricle to perfuse the lung tissue. To isolate T cells, lung tissues or mediastinal lymph nodes from 2 to 3 newborn mice were pooled together, diced, and incubated (37 °C, 30 min) with shaking (150 rpm) in digestion buffer (RPMI 1640 with 10% FBS, 15 mM HEPES, 1% penicillin/streptomycin (wt/vol) and 300 U ml^−1^ collagenase VIII) and pressed through a 100-µm nylon strainer and centrifuged (4°C, 400 x g, 5 min). The cells were resuspended in RPMI 1640 with 10% FCS. The pooled preparations constituted a single data point in our analysis.

For bone marrow analysis, we pooled freshly isolated femurs and tibias from 2-3 newborn mice. Bone marrow was released by gently crushing femurs and tibias. The resulting single-cell suspension was pressed through a 100 μm nylon strainer. The pooled preparations of bone marrow cells were a single data point in our analysis. Common lymphoid progenitors (CLP) were defined as CD45^+^ Lineage (CD3, CD4, CD11b, CD11c, CD19, F4/80, Ly6G)^-^ Sca1^lo^CD117^+^CD127^+^ α4β7^-^. Lymphoid progenitors (LP) were defined as CD45^+^ Lineage (CD3, CD4, CD11b, CD11c, CD19, F4/80, Ly6G)^-^ Sca-1^lo^ CD127^+^α4β7^+^CD117^+^ cells.

#### Murine flow cytometry

CD8^+^ T cells were isolated from digested lung tissues and mediastinal lymph nodes by negative magnetic selection with anti-CD8a microbeads (Miltenyi, Cat. No. 130-095-236) as per manufacturer instructions. After excluding dead cells, ∼10,000 cells were used for cytometry. Spleens were homogenized using a steel mesh. We used the following protocol (https://www.protocols.io/view/Cell-Surface-Immunofluorescence-Staining-Protocol-excbfiw) for our antibody staining. Cells (10^4^) were incubated (4°C, 30min) with anti-mouse CD16/CD32 to block Fc receptors. For cell surface markers and tetramers, cells were next incubated (30min, RT) with SIINFEKL (1:100), Live/Dead marker (1:1000) and antibodies [Clone IDs and dilutions listed in **table S5**]. For transcription factors (TF) and intracellular cytokines, cells were incubated (4°C, 60 min) with PMA/ionomycin and protein transport inhibitor cocktail (eBioscience, Cat. No. 4980-93), fixed (2% paraformaldehyde) and permeabilized using the True-Nuclear Transcription Factor buffer set (BioLegend, Cat. No. 424401) before adding antibodies to TFs. Stained cells were then washed (2×), resuspended in flow cytometry buffer (eBioscience, Cat. No. 4222-26) and analyzed using an Aurora CS (CYTEK). Data analyses were performed with FlowJo 10 (TreeStar).

#### Single-cell RNAseq

CD8^+^ T cells were isolated from digested lung tissues by negative magnetic selection with anti-CD8a microbeads (Miltenyi, Cat. no. 130-095-236) as per manufacturer instructions. After excluding dead cells, ∼20,000 cells were then loaded into one channel of the Chromium system using the v3 Single Cell Reagent Kit (10x Genomics) at the Cincinnati Children’s Hospital Medical Center DNA Sequencing and Genotyping Core. After capture and lysis, complementary DNA (cDNA) was synthesized and amplified as per the manufacturer’s protocol (10x Genomics). The amplified cDNA was used to construct Illumina sequencing libraries that were each sequenced using an Illumina HiSeq 4000.

Raw sequencing data were aligned to the murine reference mm10 with Cell Ranger 1.3 (10x Genomics), generating an expression count matrix. Cells that had fewer than 750 unique molecular identifiers (UMIs) or greater than 15,000 UMIs, as well as cells that contained greater than 20% of reads from mitochondrial genes or ribosomal RNA genes (RNA18S5 or RNA28S5) or hemoglobin genes, were considered low quality and were removed from further analysis. Putative multiplets were removed with DoubletFinder (version 2.0). Genes that were expressed in fewer than 10 cells were removed from the final count matrix.

#### Sequencing analysis

The Seurat package (version 3.1.0; https://satijalab.org/seurat/) was used to identify common cell types across different experimental conditions, differential expression analysis, and most visualizations. Percentages of mitochondrial genes, ribosomal genes, and hemoglobin genes were regressed during data scaling to remove unwanted variation due to cell quality using the SCTransform() function in Seurat. Principal components analysis was performed using the 3000 most highly variable genes, and the first 20 principal components (PCs) were used to perform uniform manifold approximation and projection to embed the dataset into two dimensions. Next, the first 20 PCs were used to construct a shared nearest neighbor (SNN) graph [FindNeighbors ()], and this SNN was used to cluster the dataset [FindClusters ()]. Manual annotation of cellular identity was performed by finding differentially expressed genes for each cluster using Seurat’s implementation of the Wilcoxon rank sum test [FindMarkers()] and comparing those markers to known cell type–specific genes. Global differential gene expression profiles between all cell types were identified and organized with the software cellHarmony using these Seurat parameters [fold change > 1.2, empirical Bayes *t* test *P* < 0.05, false discovery rate (FDR) corrected].

#### Automated annotation of T cell subsets

This annotation was performed using web application (https://azimuth.hubmapconsortium.org). Anchors between the reference (multimodal reference dataset of > 100,000 PBMC) and our dataset were identified using a precomputed, supervised PCA of the reference dataset that maximally captures the structure of the weighted n neighbor (WNN) graph. Cell type labels from the reference dataset were transferred to each cell of the query data set through previously identified anchors.

#### Pathway and gene ontology analysis

Differentially expressed genes between control and dysbiotic infants within with a fold change of ≥ 1.2 and t-test P-value of ≤ 0.05 (FDR corrected) were used for functional enrichment analysis of biological processes and pathways using GO-Elite. Overrepresented pathways in each subcluster were visualized with ggplot2.

#### Gene module scoring analysis

Seurat function AddModuleScore() was used to score single cells by expression of a list of genes of interest (**table S2**). This function calculates a module score by comparing the expression of an individual query gene to other randomly selected control genes expressed at similar amounts to the query genes, allowing a robust method for scoring modules containing both lowly and highly expressed genes and scoring cells with different sequencing depths.

#### Identification of transcription factors

We predicted transcription factors regulating T cell heterogeneity using SCENIC (single-cell regulatory network inference and clustering). Default parameters were used for the SCENIC workflow in R and the normalized single-cell gene expression matrix for T cell clusters from Seurat was used as input. Co-expression analysis was performed with GENIE. For visualization, we calculated the average regulon activity (AUC) scores for each CD8 T cell cluster and selected the top regulons to plot as a heatmap using R package “pheatmap”.

#### NFIL3-Chromatin Immunoprecipitation and sequencing (ChipSeq)

Briefly, 10^5^ CD8^+^ T cells were fixed with 1% formaldehyde, processed using truChIP Chromatin Shearing Reagent Kit (Covaris), and sonicated using Covaris S2 ultrasonicator. Sheared chromatin was immunoprecipitated with an anti-NFIL3 antibody (Clone PM097, MBL Bio). DNA segments were end-repaired and ligated to indexed Illumina adaptors using TruSeq ChIP Library Preparation Kit (Illumina). The resulting libraries were sequenced with the Illumina HiSeq2000. ChIP-seq reads were trimmed using Trimmomatic (version 0.38) to remove primer and low-quality bases. Reads with mapping quality below 30 were discarded and then aligned to murine reference (mm10) using bowtie2 (version 2.5.0), using a maximum insert size of 2000 bp, with the “--no-mixed” and “--no-discordant” parameters. Duplicate reads and reads mapping to the blacklist regions and mitochondrial DNA were removed. ChIP-seq peaks were called in each replicate versus an input IgG control, using MACS2 (version 2.2.7.1) with a q-value cutoff of 0.001, and overlapping peaks among replicates were merged. Diffbind (version 2.10.0) was used to identify differentially abundant peaks between control and dysbiotic CD8^+^ T cells. For motif analysis of the identified NFIL3 peaks, the sequences of ±100 bps flanking the peak summits were used as input to findMotifsGenome.pl from HOMER for de novo motif discovery.

#### ChIP-qPCR for Tcf7 and Lef1

Approximately, 10^4^ CD8^+^ T cells isolated from digested lung tissues by negative magnetic selection with anti-CD8a microbeads were fixed (10 min, RT) in 1% formaldehyde, quenched (10 min, RT) with 125 mM glycine and washed with TE 0.1% SDS with protease inhibitors and sonicated using an S220 Focused-ultrasonicator (Covaris). Prior to immunoprecipitation, sheared chromatin was precleared (20 min, 4°C) using Protein G Dynabeads (Thermo Fisher Scientific). Immunoprecipitations were performed using fresh beads and anti-Histone H3K4me3 antibody (#07-473, Millipore) using a SX-8G IP-STAR automated system (Diagenode) with the following wash buffers: (1) RIPA 150 mM NaCl, (2) RIPA 400 mM NaCl, (3) Sarkosyl Buffer (2 mM EDTA, 50 mM Tris-HCl, 0.2% Lauroylsarcosine sodium salt, and (4) TE 0.2% Triton X-100. Immunoprecipitated chromatin was treated with Proteinase K (Thermo Fisher Scientific) (42°C, 30 min, 65°C, 4 hr, and 15°C,10 min) in elution buffer (TE 250 mM NaCl 0.3% SDS). Phenol: chloroform isoamyl alcohol with Tris-HCl (pH 8.0) and chloroform phase separation were used to isolate DNA, followed by overnight ethanol precipitation. ChIP DNA was treated with molecular grade RNase and subjected to quantitative real-time PCR using the custom-made primer pairs listed in **Table S3**. Data were analyzed as percent enrichment over input DNA.

#### Tcf7 Gene expression

RNA from 10^4^ CD8^+^ T cells was isolated using the RNeasy Kit (Qiagen) following the manufacturer’s protocol. cDNA was synthesized using the Verso reverse transcriptase kit (Thermo Fisher) following the manufacturer’s protocol. Real-time PCR was performed using SYBR green (Applied Biosystems) and analyzed using primer sets (table S3)

#### Human Studies

Human tissues were obtained from deceased subjects through an approved protocol at CCHMC. The study does not qualify as human subjects research as determined by CCHMC IRB because tissue samples were obtained from deceased individuals. Lung tissues were processed for single-cell suspensions, as previously described(*11*). Briefly, lung tissue was minced into 2 mm pieces and incubated (RT, 30 min) with a prewarmed digestion solution of collagenase II (0.2 mg ml^−1^) and DNase I (0.1 mg ml^−1^) in PneumaCult-EX with gentle mechanical agitation. Dissociated single lung cells (10^5^ cells) were labeled for 30 min with streptavidin-coupled anti-CD3 Ab (Cat. No 480133, BioLegend). The magnetically labeled fraction was retained using a magnetic separator (MACS) with magnetic streptavidin beads according to the manufacturer’s protocol and used for single-cell RNAseq analysis and flow cytometry.

#### Single cell RNAseq and analysis from human newborn lungs

After excluding dead cells, ∼5,000 cells per sample were then loaded into one channel of the Chromium system using the v3 single-cell reagent kit (10X Genomics). Following capture and lysis, cDNA was synthesized and amplified as per the manufacturer’s protocol (10X Genomics). The amplified cDNA was used to construct Illumina sequencing libraries that were each sequenced using an Illumina HiSeq 4000. Raw sequencing data were aligned to the GRCh38 reference genome, generating expression count matrix files. Cells that had fewer than 750 unique molecular identifiers (UMIs) or greater than 15,000 UMIs, as well as cells that contained greater than 20% of reads from mitochondrial genes or rRNA genes (RNA18S5 or RNA28S5) or hemoglobin genes, were considered low quality and removed from further analysis. Putative multiplets were removed with DoubletFinder (version 2.0). Genes that were expressed in fewer than 10 cells were removed from the final count matrix. The Seurat package (version 4.1.0, https://satijalab.org/seurat/) was used to identify common cell types across different experimental conditions, differential expression analysis, and most visualizations. Percentages of mitochondrial, ribosomal genes, and hemoglobin genes were regressed during data scaling to remove unwanted variation due to cell quality using the SCTransform() function in Seurat. PCA was performed using the 3,000 most highly variable genes, and the first 20 principal components (PCs) were used to perform UMAP to embed the dataset into two dimensions. Next, the first 20 PCs were used to construct a shared nearest neighbor graph (SNN; FindNeighbors ()) and this SNN used to cluster the dataset (FindClusters ()). Manual annotation of cellular identity was performed by finding differentially expressed genes for each cluster using Seurat’s implementation of the Wilcoxon rank-sum test (FindMarkers()) and comparing those markers to known cell type-specific genes from published studies (*12*). We used DESeq2, for analysis of aggregated read counts. Briefly, for cells of a given type, we first aggregated reads across biological replicates, transforming a genes-by-cells matrix to a genes-by-replicates matrix using matrix multiplication. We used both a Wald test of the negative binomial model coefficients (DESeq2-Wald) to compute the statistical significance (fold change > 1.2, empirical Bayes t-test *P*-value <0.05, FDR corrected).

#### Assessment of effector function by flow cytometry

Briefly, 1 × 10^4^ CD3ε^+^ cells from the lung were labeled for 30 min with PE-coupled peptide-loaded Influenza-specific HLA-A*02/M1_58– 66_(GILGFVFTL) tetramers (NIH Tetramer Core facility). Virus-specific CD8^+^ T cells were magnetically (MACS) enriched using anti-phycoerythrin (PE) magnetic beads (Miltenyi) according to the manufacturer’s protocol. Enriched influenza-specific-specific CD8^+^ T cells were used for multiparametric flow cytometry analysis.

#### For in vitro expansion of virus-specific human CD8^+^ T cells

1 × 10^4^ CD3ε^+^ cells from the lung were stimulated with influenza-specific HLA-A*02/M1_58–66_(GILGFVFTL) tetramers (5 µM) and beads coated with anti-CD3/anti-CD28/anti-CD2 mAb (all 0.5 µg ml^−1^) and expanded for 14 days in a complete RPMI culture medium containing recombinant IL2 (20 IU ml^−1^). Cells were incubated (4°C, 30min) with Human Fc Block (BD Bioscience, Cat. No. 564219). Next, cells were next incubated (30min, RT) with GILGFVFTL (1:100), 7-AAD (Cat. No. 559925; dilution 1:33), and a cocktail of anti-CD4 (Cat. No. L200, dilution 1:200), anti-CD8 (Cat. No. SK1, dilution 1:100), anti-CD27 (Cat. No. L128, dilution 1:200), anti-CD28 (Cat. No. CD28.2, dilution 1:100), anti-CD45RA (Cat. No. HI100, dilution 1:800), anti-CD69 (Cat. No. FN50, dilution 1:50), anti-CD45RA (Cat. No. HI100, dilution 1:200), anti-CD25 (Cat. No. BC96, dilution 1:33), anti-CD103 (Cat. No. No. 350215, dilution 1:50), anti-Ki67 (Cat. No. No. 350513, dilution 1:100) (all from Biolegend), anti-TCF-1 (Cat. No. C63D9, dilution 1:100) (from Cell signaling) and anti-KLRG-1 (Cat. No. 13F12F2, dilution 1:50) (from Biosciences). FoxP3/transcription factor staining buffer and fixation/permeabilization solution kit (Cat. No. 562765, BD Biosciences) were used according to the manufacturers’ instructions to stain with anti-IFN (BD Biosciences, Cat. no. 25723.11, dilution 1:8). Stained cells were fixed in paraformaldehyde (2%) (Sigma) and analyzed using Aurora CS (CYTEK). Data analyses were performed with FlowJo 10 (TreeStar).

#### Fecal microbiome sequencing and analysis from human newborns

Genomic DNA extracted from matched frozen stool specimens using the Fecal Microbiome DNA Kit (Promega) was used to prepare DNA libraries with the Illumina Nextera DNA Flex library kit (Illumina, San Diego, CA). Pooled libraries were sequenced using Illumina NovaSeq. FASTQ files were quality-filtered and screened to remove human sequences using Kneaddata(*13*). Metagenomic samples were identified using MetaPhlAn2(*14*), which uses a library of clade-specific markers to provide pan-microbial (bacterial, archaeal, viral, and eukaryotic) profiling (with default settings) in combination with HUMAnN2(*15*) for functional metagenomic profiling(*16*) (with default settings). MelonnPan was used to predict metabolite compositions from the metagenomic sequences.

#### Untargeted Metabolomic assessment

Metabolites in serum were extracted in 50% methanol, centrifuged, and the resulting supernatants were diluted 1:20. Ultra-high performance liquid chromatography mass spectrometry (UHPLC-MS) data were then acquired on a Q Exactive™ HF Mass Spectrometer in negative ion full scan mode (50-750m/z). Metabolites were separated via UHPLC using a binary solvent mixture of 20mM ammonium formate at pH 3.0 in LC-MS grade water and 0.1% formic acid (%v/v) in LC-MS grade acetonitrile in conjunction with a Syncronis™ column. For all runs the sample injection volume was 2uL. Metabolite data were analyzed using the XCMS using default parameters (*17*). Metabolites were identified by matching observed m/z signals (+/-10ppm) and chromatographic retention times to those observed from commercial metabolite standards (Sigma).

#### Statistical analysis

All data met the assumptions of the statistical tests used. Statistical tests used for microbiome or single-cell analyses are described in relevant sections. For comparing the differences between groups, we used either unpaired two-tailed Student’s *t*-test or analysis of variance (ANOVA) or Wilcoxon signed-rank test. We used the Pearson correlation coefficient to measure the correlation between different variables. We used the Kaplan-Meier log-rank test to compare morbidity between groups (All in GraphPad Prism 10). *P*-values are indicated as follows: * *P* ≤ 0.01.

